# Establishment of a green fluorescent protein (GFP)-based reporter for picornaviral 3C proteases

**DOI:** 10.1101/2025.07.31.667855

**Authors:** Junki Hirano, Tsuyoshi Hayashi, Yuichi Someya, Kazuma Okada, Kentaro Uemura, Ming Te Yeh, Chikako Ono, Shuhei Taguwa, Yoshiharu Matsuura

**Author notes:** Corresponding author: Yoshiharu Matsuura, DVM, PhD Laboratory of Virus Control Center for Infectious Disease Education and Research, and Research Institute for Microbial Diseases, University of Osaka, 1-10, Yamada-oka, Suita, Osaka 565-0871, Japan.

## Abstract

Picornaviruses represent a diverse group of plus-stranded RNA viruses, many of which have been linked to severe diseases in both humans and animals. The viral 3C protease is essential for the maturation of viral proteins and the propagation of picornaviruses and, owing to its cleavage activity against multiple host proteins, is associated with the pathogenesis of picornaviruses. The picornaviral 3C protease is an ideal drug target for inhibiting viral propagation and mitigating pathogenesis; however, methodology to evaluate and compare the activity of phylogenetically diverse proteases remains lacking. To address this, herein, we propose a novel green fluorescent protein (GFP)-based reporter optimized to visualize the enzymatic activity of picornaviral 3C proteases in cells by using the conformational change of a GFP variant induced by the 3C protease, generating fluorescence emission linked to the enzymatic activity. Upon treatment of picornaviruses with a known 3C protease inhibitor, the fluorescence decreased in a dose-dependent manner, demonstrating that the signal depended on the activity of the 3C protease. The reporter system for the 3C protease can be applied to major pathogenic human picornaviruses, such as those in the genera *Enterovirus*, *Rhinovirus*, *Cosavirus*, *Salivirus,* and *Kobuvirus*. Furthermore, the fluorescent signal from the reporter was confirmed in various animal-derived picornaviruses, such as those from bats, rodents, and primates. Therefore, the reporter could be widely used to analyze the activity of several 3C proteases from currently prevalent picornaviruses and those that may emerge in the future. To demonstrate the flexibility of the reporter in comparing phylogenetically different proteases, the enzymatic activity of the 3C protease derived from clinical strains of enterovirus A71 (EV-A71) was tested and compared. The results showed that the amino acid residues of the 3C protease affect its activity by utilizing the reporter system. Additionally, clinical EV- A71 strains had different effects on the activity of 3C protease against host proteins. Our findings will aid in elucidating the molecular characteristics of 3C proteases among picornaviruses and developing therapeutics to mitigate the pathogenesis of these viruses.

## Importance

The family Picornaviridae comprises various biologically distinct viruses that have been linked to life-threatening diseases, including emerging and previously neglected pathogens whose molecular characteristics remain elusive. Although the viral protease encoded by picornaviruses, the 3C protease, is an ideal target for anti-picornaviral pharmacological intervention, the current assay system for evaluating the activity of 3C proteases remains limited to a few specific picornaviruses. To address this, we sought to establish a flexible and straightforward protocol for assessing and comparing various picornaviral 3C proteases. We developed a cell-based assay for 3C proteases using a split-green fluorescent protein (GFP) variant with a specific viral sequence that could be applied to a wide range of human pathogenic and animal-derived picornavirus strains. Furthermore, by swapping the viral sequence inside GFP for a host-derived sequence, this assay could be used to evaluate the 3C protease activity of cleaving host proteins. This assay will facilitate future research comparing functional differences of 3C proteases among picornaviruses, conducting high-throughput screening of anti-picornaviral drugs, or investigating the relationship between protease activity and picornaviral pathogenesis.

## Introduction

The family Picornaviridae comprises more than 200 viral serotypes with highly diverse characteristics (1). Human pathogenic picornaviruses, such as viruses belonging to the genera *Enterovirus*, *Rhinovirus*, *Hepatovirus*, *Parechovirus*, and *Kobuvirus*, have been linked to various symptoms, including poliomyelitis, aseptic meningitis, herpangina, hand-foot-and-mouth disease (HFMD), myocarditis, acute hepatitis, and acute gastroenteritis (2, 3). Despite the significant risk posed by picornavirus infections on global public health, effective preventive measures or therapeutics against them are lacking, except for potent vaccines against polioviruses (4–7). Moreover, several novel picornaviruses have been identified in animal specimens. Owing to, among others, the lack of robust cell culture systems, the pathogenicity and natural host range of animal picornaviruses are mostly undetermined; however, these viruses have been detected in zoonotic reservoirs and humans (8–12). The molecular characteristics of these animal viruses remain to be fully elucidated.

*Enterovirus* A71 (EV-A71), a picornavirus of the genus *Enterovirus* that causes febrile illnesses and HFMD in infants (13), circulates in the Asia-Pacific region and has been responsible for severe outbreaks of HFMD that are occasionally accompanied by encephalopathy, encephalitis, and acute flaccid paralysis, among other severe neurological disorders (14). Although the transmission and circulation of EV-A71 among the population creates polymorphisms within the viral genome, the molecular determinants within the genome that influence virulence have not been fully elucidated. This virus harbors a single-stranded, positive RNA genome. Following the attachment and entry of EV-A71 particles, viral RNA is released into the cytoplasm and subsequently translated into a large single polyprotein (∼ 2,200 amino acids). The polyprotein is proteolytically processed by viral 2A and 3C proteases to produce four structural, namely VP1, VP2, VP3, and VP4, and seven nonstructural, namely 2A, 2 B, 2C, 3A, 3 B, 3C, and 3D, proteins (15). 2A protease cleaves between the structural and nonstructural regions of the polyprotein, whereas 3C protease cleaves other regions and plays an indispensable role in viral propagation (16). The viral life cycle completely depends on the activity of the 3C protease, making it an attractive target for pharmacological intervention. Furthermore, the 3C protease cleaves multiple host proteins, such as CSTF2, eIF5b, G3BP1, IRF7, NLRP1, and TBP (17–24). Therefore, determining how these cleavage events orchestrate viral pathogenesis is necessary to gain insights into enterovirus-induced diseases.

Recently, a robust fluorogenic system has been implemented to visualize the cleavage activity of proteases inside cells (25). The system with a large dynamic range designed split-green fluorescent protein (GFP) to induce a conformational change and reconstitution of active GFP depending on the activity of the protease of interest (Fig. 1A). The fluorogenic reporter fixed the 10^th^ and 11^th^ beta-strands of GFP (GFPβ10 and GFPβ11) with a linker containing cleavage sequence of the protease and arranged the structure incapable of interacting with the rest of the GFP beta-barrel (GFPβ1-9). In the presence of the protease, the target sequence is cleaved, inducing a conformational change of GFPβ10 and GFPβ11, thereby allowing access to GFPβ1-9. This protease reporter, named flipGFP, has been used to visualize *in vivo* caspase activity in zebrafish and *Drosophila* (25).

**Fig. 1.**
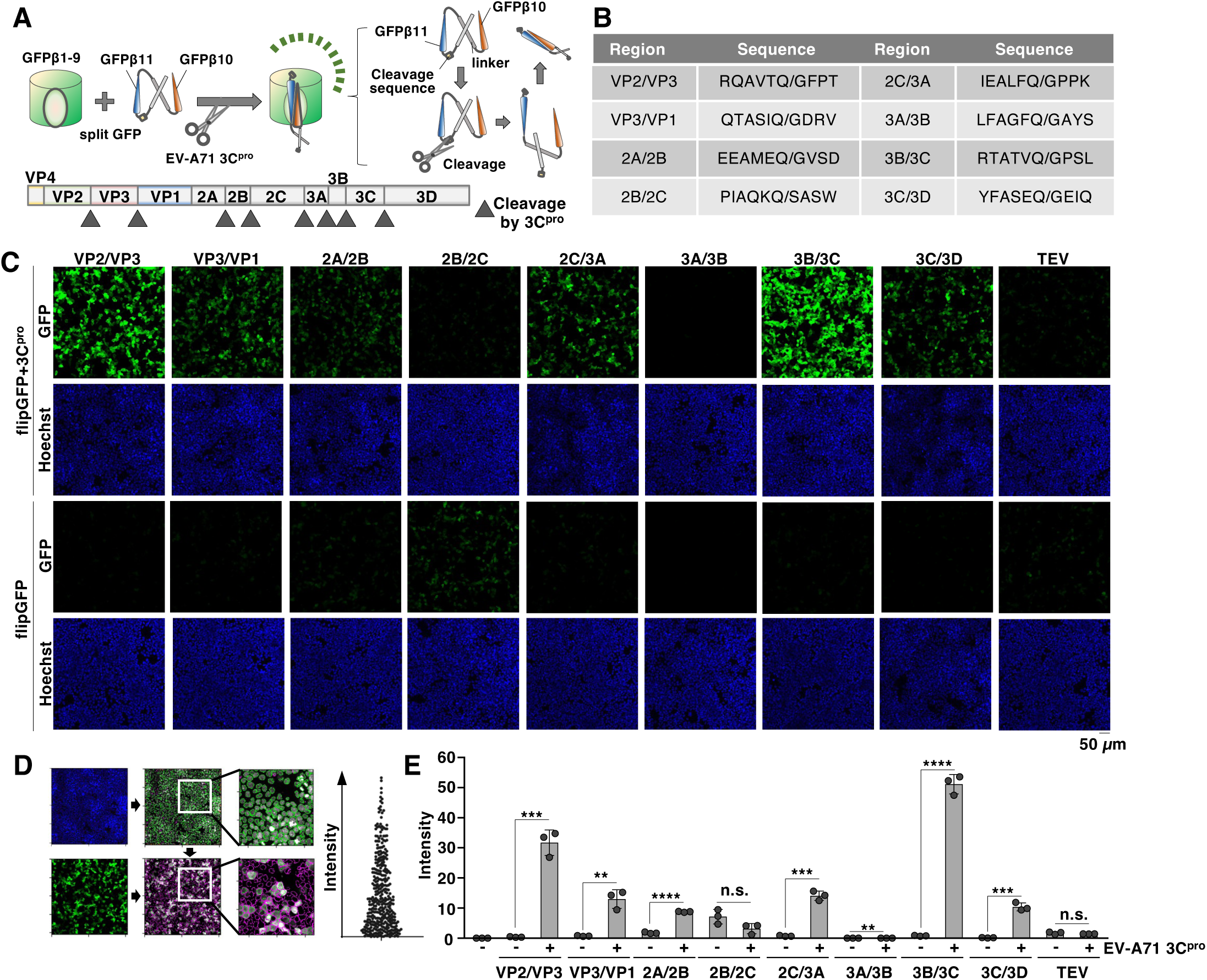
Optimization of cleavage sequence inside flipGFP for EV-A71 3C protease. (A) Schematic representation of the flipGFP system. The cleavage sequence of the protease is inserted between the 10^th^ and 11^th^ beta-strands of GFP (GFPβ10 and GFPβ11) of split GFP. In the presence of protease, the sequence was cleaved by the protease, inducing the conformational change of the split GFP. The changed structure of split GFP interacts with the rest of the GFP beta-barrel (GFPβ1-9) (*Upper*). A schematic representation of the genomic structure of EV-A71 is also shown. The EV-A71 RNA is translated into a single polyprotein that is subsequently processed into viral proteins through cleavage by viral 3C proteases at the site indicated by the arrowhead (*Lower*). (B) The cleavage region inside the EV-A71 polyprotein and the corresponding amino acid sequence inserted into flipGFP. (C) HEK293T cells were transfected with the plasmid encoding 3C protease and the indicated flipGFP. The fluorescent signal of flipGFP was monitored using fluorescent microscopy. (D) Schematic representation of the quantification of GFP-positive cells. Each cell visualized using the Hoechst 33342 staining was analyzed for the intensity of fluorescence using the CellProfiler software (Cimini Lab, the Broad Institute of MIT and Harvard) (*Left*). The fluorescent intensity of each cell inside the captured images was calculated. The average intensity was used for the following studies (*Right*). (E) HEK293T cells were transfected with the plasmid encoding 3C protease and the indicated flipGFP. The fluorescent intensity was calculated 24 h post-transfection. The data presented in C and E are representative of two independent experiments. For the experiment presented in E, significance was determined using Student’s *t-*test (n = 3) (**P ≤ 0.01; ***P ≤ 0.001; ****P ≤ 0.00001; n.s., not significant).

Although individual reporters monitoring the activity of 3C proteases derived from a few picornaviruses have been reported, especially in the genus *Enterovirus* (26, 27), an unbiased protease reporter for the diverse Picornaviridae family remains lacking. Moreover, existing studies have failed to characterize the polymorphism of the 3C protease within specific viral species due to the obstacle associated with establishing multiple assays optimized for each protease. Therefore, in this study, we aimed to develop and validate a flipGFP optimized to monitor the activity of a wide range of picornaviral 3C proteases. Furthermore, using EV-A71 as a model, we elucidated the mechanism by which 3C protease polymorphisms affect the cleavage activity of host proteins.

## Results

### Establishment of a flipGFP-based reporter for EV-A71 3C protease

To assess the activity of the EV-A71 3C protease, we initially examined the sequence within flipGFP cleaved by the protease and constructed expression vectors for flipGFP encoding a known cleavage sequence inside the EV-A71 polyprotein (e.g., the sequence derived from the VP2/VP3, VP3/VP1, 2A/2B, 2B/2C, 2C/3A, 3A/3B, 3B/3C, or 3C/3D region) (Fig. 1B). The fluorescent signal of GFP was observed in HEK293T cells co-expressing the 3C protease with flipGFP encoding the VP2/VP3, VP3/VP1, 2A/2B, 2C/3A, 3B/3C, or 3C/3D sequences, but not in those co-expressing the 2B/2C, 3A/3B, or control TEV protease cleavage sequence (ENLYFQG). To quantify the observed GFP signal, the signal intensity of each cell was calculated and normalized using the optimized pipeline of CellProfiler software (Fig. 1D). Analysis of the fluorescent signal of each flipGFP showed significant signal intensity, especially in the VP2/VP3 or 3B/3C sequences of flipGFP (Fig. 1E). Therefore, we explored the potential of these flipGFPs to monitor 3C protease activity.

To determine whether the fluorescence observed in HEK293T cells expressing flipGFP depended on the enzymatic activity of 3C or the precursor 3CD protease, we generated inactive mutants of these proteases by substituting cysteine 147 with serine (C147S) (28) (Fig. 2A). Inactive 3C or 3CD proteases showed undetectable fluorescence signals when co-expressed with flipGFP (3B/3C) (Fig. 2B and C) or flipGFP (VP2/VP3) (Fig. 2D and E). In addition, treatment with rupintrivir, a known inhibitor of 3C protease, dose-dependently reduced the fluorescence signal in HEK293T cells co-expressing 3C protease and flipGFP (3B/3C) (Fig. 2F and G) or flipGFP (VP2/VP3) (Fig. 2H and I).

**Fig. 2.**
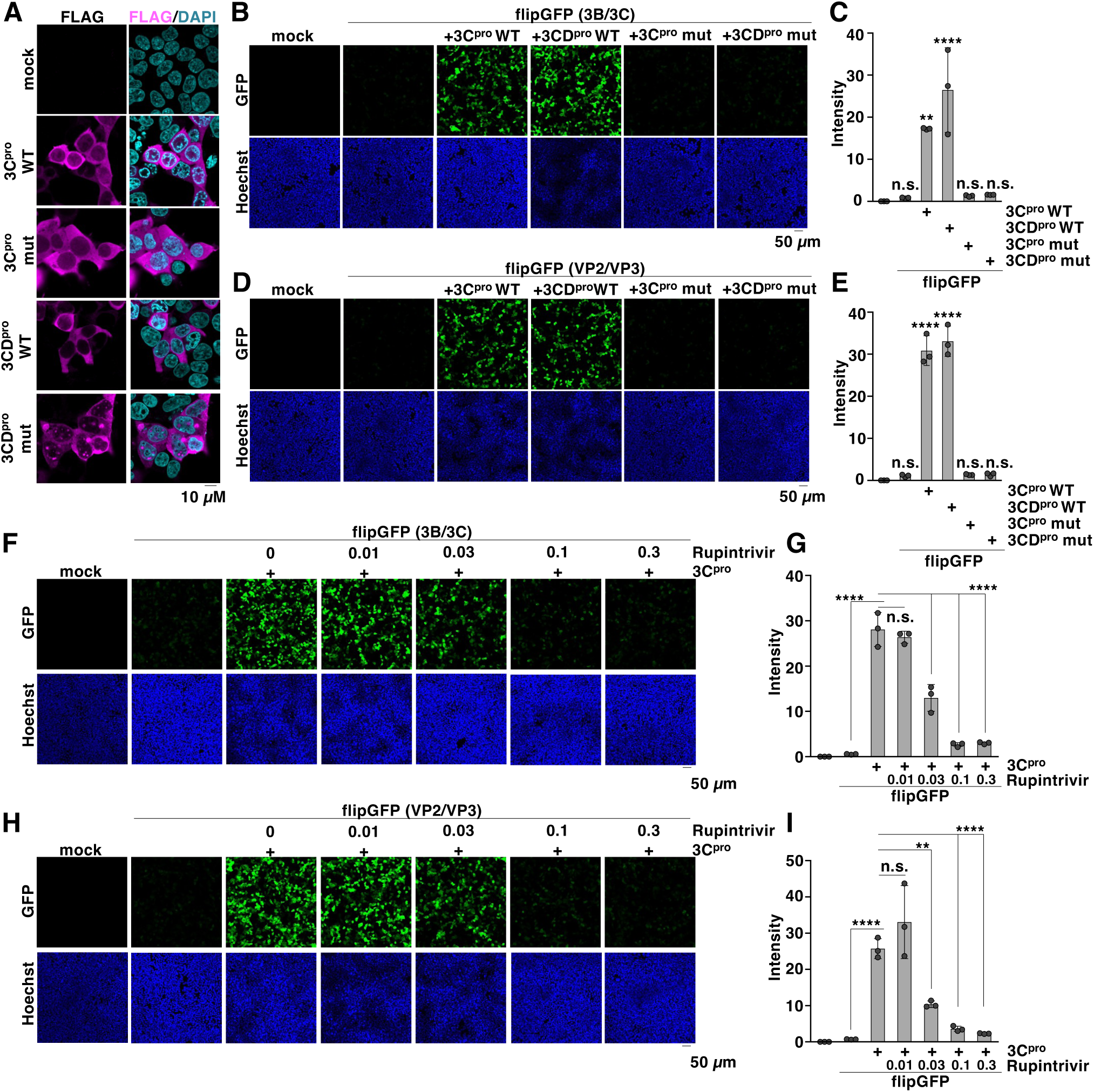
Fluorescence of flipGFP depends on the activity of 3C protease. (A) Wild-type or mutant 3C or 3CD protease was expressed in HEK293T cells. The expression of each 3C protease was detected with fluorescent microscopy using a FLAG antibody. (B) Wild-type or mutant 3C or 3CD protease was transfected along with flipGFP (3B/3C). The fluorescent signals were detected using microscopy at 24 h post-transfection. (C) Wilde-type or mutant 3C or 3CD protease was transfected along with flipGFP (3B/3C). The fluorescent intensity was calculated 24 h post-transfection. (D) Wild-type or mutant 3C or 3CD protease was transfected along with flipGFP (VP2/VP3). The fluorescent signals were detected using microscopy at 24 h post-transfection. (E) Wild-type or mutant 3C or 3CD protease was transfected along with flipGFP (VP2/VP3). The fluorescent intensity was calculated 24 h post-transfection. (F) HEK293T cells transfected with the expression vector of 3C protease and flipGFP (3B/3C) were treated with rupintrivir at concentrations of 0.01, 0.03, 0.1, or 0.3 µM. The fluorescent signals were detected using microscopy at 24 h post-transfection. (G) HEK293T cells transfected with the expression vector of 3C protease and flipGFP (3B/3C) were treated with rupintrivir at concentrations of 0.01, 0.03, 0.1, or 0.3 µM. The fluorescent intensity was calculated 24 h post-transfection. (H) HEK293T cells transfected with the expression vector of 3C protease and flipGFP (VP2/VP3) were treated with rupintrivir at concentrations of 0.01, 0.03, 0.1, or 0.3 µM. The fluorescent signals were detected using microscopy at 24 h post-transfection. (I) HEK293T cells transfected with the expression vector of 3C protease and flipGFP (VP2/VP3) were treated with rupintrivir at concentrations of 0.01, 0.03, 0.1, or 0.3 µM. The fluorescent intensity was calculated 24 h post-transfection. The data presented in A-I are representative of two independent experiments. For the experiments presented in C, E, G, and I, significance was determined using a one-way ANOVA test (n = 3) (**P ≤ 0.01; ****P ≤ 0.00001; n.s., not significant).

Collectively, these data suggest that the observed signal from flipGFP carrying 10 amino acid residues in the target sequence of EV-A71 3C protease (sequence 3B/3C or VP2/VP3) corresponds to the activity of the 3C protease inside the cells.

### flipGFP-based reporter could detect the activity of 3C protease from diverse picornaviruses

The family Picornaviridae comprises diverse categories of viruses with genetically distinct features (Fig. 3A). To further explore the applications of flipGFP, we established a universal system for assessing 3C proteases derived from a variety of Picornaviridae family members. We constructed the vector expressing 3C protease of the genus *Enterovirus* (coxsackievirus A6 (CV-A6), coxsackievirus A16 (CV-A16), coxsackievirus B3 (CV-B3), echovirus 18 (echovirus), poliovirus 1 (PV1), and enterovirus D68 (EV-D68)), genus *Rhinovirus* (human rhinovirus A (HRV A), human rhinovirus B (HRV B) and human rhinovirus C (HRV C)), genus *Cosavirus* (cosavirus A (CosaV)) genus *Salivirus* (salivirus A (SaliV)) and genus *Kobuvirus* (Aichivirus). While we prepared a flipGFP construct containing the target sequence of each virus, the sequence corresponding to the VP2/VP3 region was only applicable to EV-A71 and EV-D68, but not to CVB3 or PV1 (Fig. S1A and B). In contrast, the flipGFP-inserted 3B/3C sequence could be applied to all viruses (Fig. S1C). Therefore, we primarily used the 3B/3C sequence to generate Picornaviridae flipGFP (Fig. 3C). We co-expressed Picornaviridae flipGFP with each 3C protease in HEK293T cells and observed significant upregulation of the fluorescent signal compared to the expression of flipGFP alone (Fig. 3D and E). These data suggest that the established flipGFP system can be widely used to monitor the activity of viruses belonging to the family Picornaviridae.

**Fig. 3.**
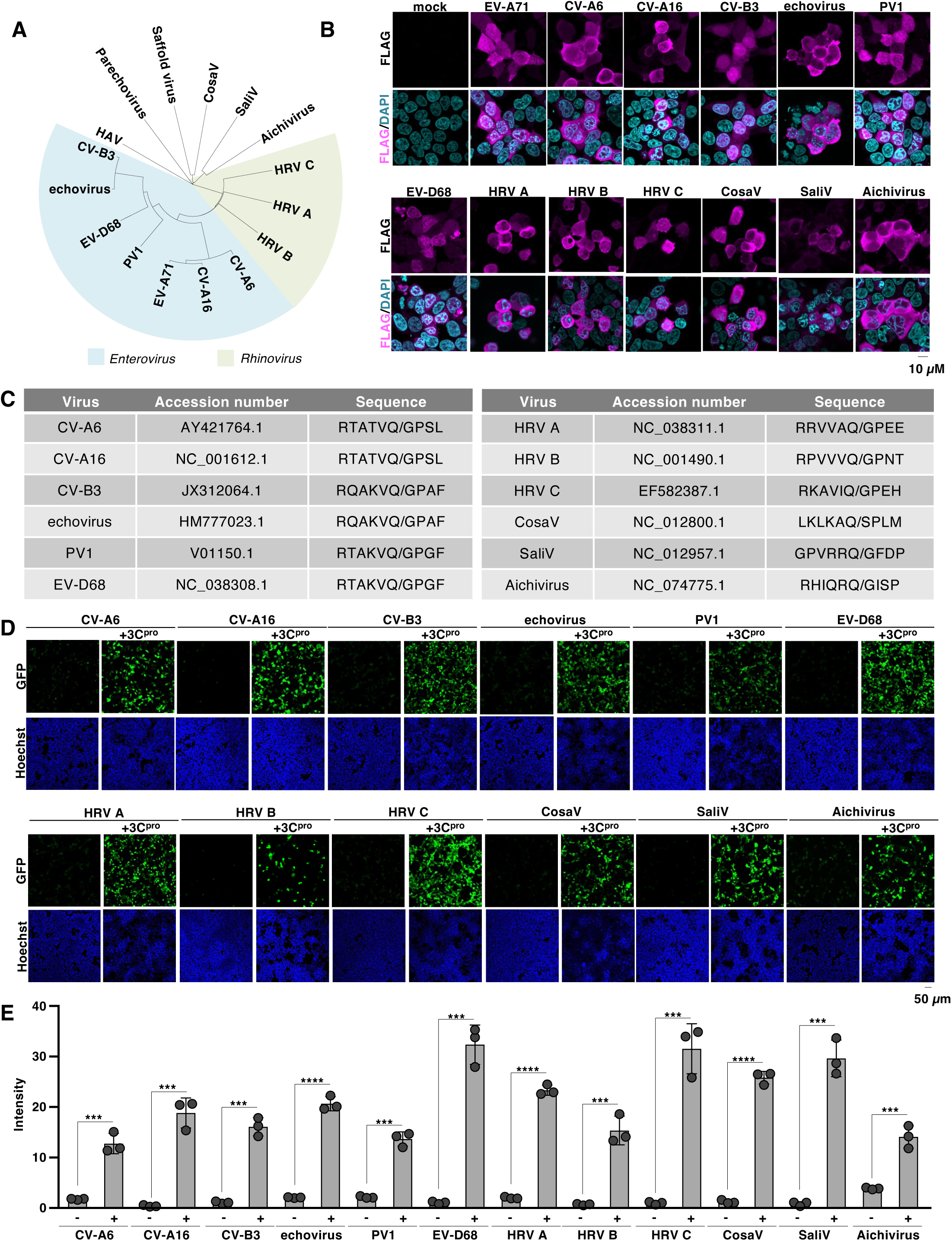
Establishment of a flipGFP-based reporter for picornaviral 3C proteases. (A) Phylogenetic tree of 3C protease derived from picornaviruses (i.e., EV-A71, coxsackievirus A6 (CV-A6), coxsackievirus A16 (CV-A16), coxsackievirus B3 (CV-B3), echovirus 18 (Echovirus), Poliovirus 1 (PV1), enterovirus D68 (EV-D68), human rhinovirus A (HRV A), human rhinovirus B (HRV B), human rhinovirus C (HRV C), hepatitis A virus (HAV), parechovirus, saffold virus, cosavirus (CosaV), sali virus (SaliV) or (Aichivirus). The amino acid sequence of each picornaviral 3C protease was aligned by Clustal Omega (EMBL-EBI) and indicated using INTERACTIVE TREE OF LIFE v7 (61). (B) The FLAG-tagged picornaviral 3C protease was expressed in HEK293T cells. The expression of each 3C protease was detected with fluorescent microscopy using a FLAG antibody. (C) The accession number of the picornavirus used in this study and the corresponding amino acid sequence between the 3B and 3C regions. (D) HEK293T cells were transfected with the plasmid encoding picornaviral 3C protease and the flipGFP optimized for each 3C protease. The fluorescent signal of flipGFP was monitored with fluorescent microscopy. (E) HEK293T cells were transfected with the plasmid picornaviral 3C protease and the flipGFP optimized for each 3C protease. The fluorescent intensity was calculated 24 h post-transfection. The data presented in B, D, and E are representative of two independent experiments. For the experiment presented in E, significance was determined using Student’s *t*-test (n = 3) (***P ≤ 0.001; ****P ≤ 0.00001).

### flipGFP could apply to novel picornaviruses detected from the reservoir of zoonosis

Developing a strategy for evaluating the protease activity of animal picornaviruses can provide deeper insights into their characteristics. We established a flip-GFP system for animal picornaviruses. Three bats (*Miniopterus pusillus*, *Miniopterus magnate,* and *Rhinolophus sinicus*), two simians (*Papio anubis* and *Macaca mulatta*), and two rodents (*Caryomys eva* and *Niviventer niviventer*) were selected for this study (8–10) (Fig. 4A and B). Sequence logo analysis of the 3B/3C cleavage sequence showed similarities between animal and human picornaviruses (Fig. 4C and D). Therefore, we generated expression vectors for the 3C protease derived from each virus, and the corresponding flipGFP- containing 3B/3C cleavage sequence. Fluorescence analysis of HEK293T cells expressing each 3C protease confirmed the cytoplasmic distribution of the protease (Fig. 4E), similar to that in human picornaviruses (Fig. 3B). Subsequently, HEK293T cells were co-transfected with a plasmid encoding animal picornavirus flipGFP and 3C protease. Significant fluorescent signals were detected in all examined flipGFPs co-expressed with the protease (Fig. 4F and G), suggesting that the flipGFP system can detect the protease activity of the animal picornaviruses. Next, to examine the effect of a known 3C protease inhibitor against animal picornaviruses, HEK293T cells co-expressing flipGFP and the picornaviral 3C protease were treated with rupintrivir. The results showed that the fluorescent signal was dose-dependently reduced following treatment, whereas its efficacy varied among the viruses (Fig. 4H and S2A-G), suggesting phenotypic differences among these proteases.

**Fig. 4.**
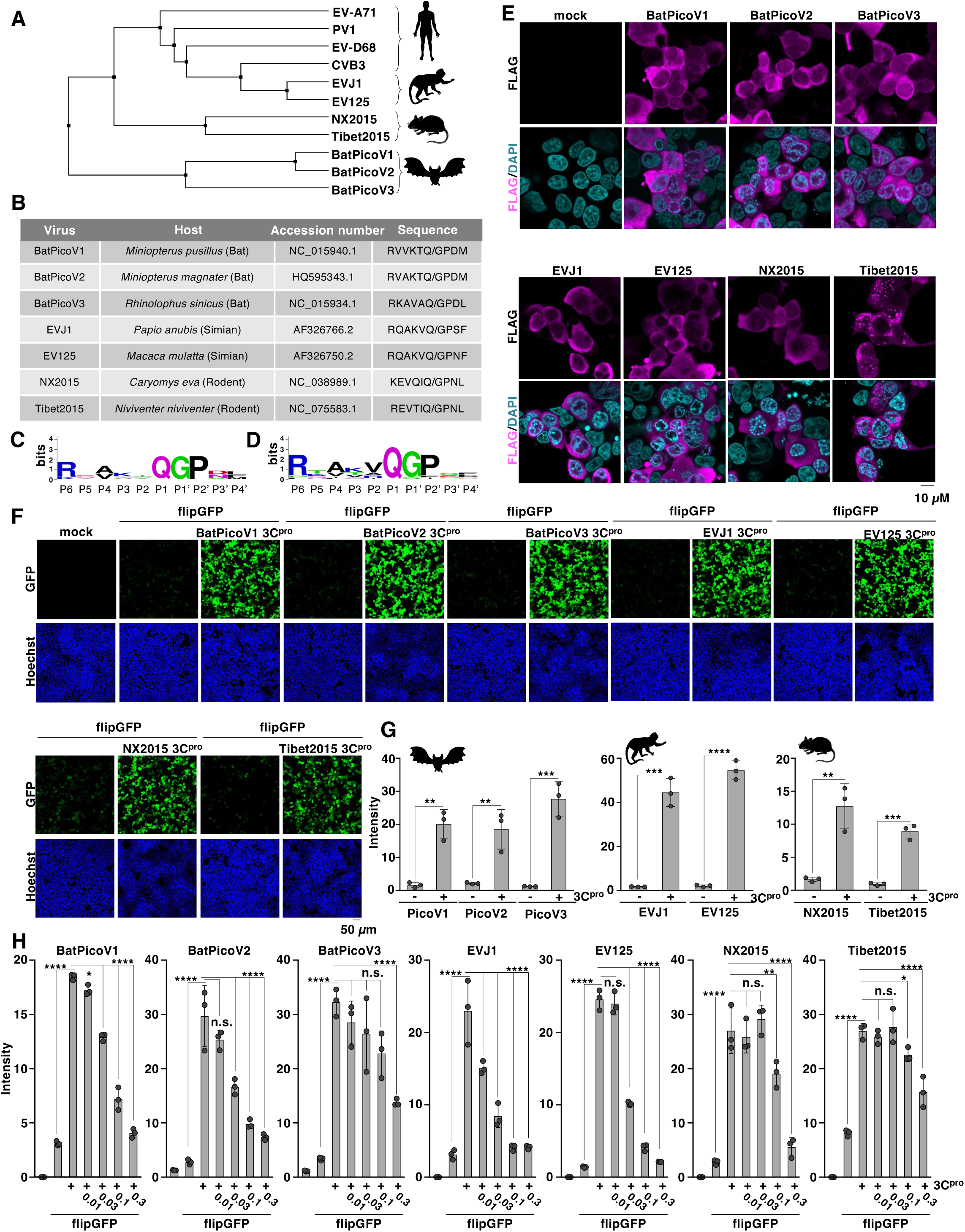
Development of a flipGFP-based reporter for 3C proteases derived from animal- **picornaviruses.** (A) Phylogenetic tree of 3C protease derived from human and animal picornaviruses (i.e., human (EV- A71, CVB3, PV1, or EV-D68), bats (BatPicoV1, BatPicoV2, or BatPicoV3), primates (EVJ1 or EV125), or rodents (NX2015 or Tibet2015). The amino acid sequence of each picornaviral 3C protease was aligned by Clustal Omega (EMBL-EBI) and indicated using Jalview software version 2.11.4.1 (62). (B) The accession number of the picornavirus used in this study, the information on the reservoir, and the corresponding amino acid sequence between the 3B and 3C regions. (C) The protease cleavage sequence presented in Figure 4B (animal picornaviruses) was analyzed and visualized using WebLogo Version 2.8.2. (D) The protease cleavage sequence presented in Fig. 3C (human picornaviruses) was analyzed and visualized using WebLogo Version 2.8.2. (E) The FLAG-tagged picornaviral 3C proteases were expressed in HEK293T cells. The expression of each 3C protease was detected with fluorescent microscopy using a FLAG antibody. (F) HEK293T cells were transfected with the plasmid encoding 3C protease and the flipGFP optimized for each 3C protease. The fluorescent signal of flipGFP was monitored with fluorescent microscopy. (G) HEK293T cells were transfected with the plasmid encoding 3C protease derived from bats, primates, or rodents, and the flipGFP optimized for each 3C protease. The fluorescent intensity was calculated 24 h post-transfection. (H) HEK293T cells transfected with the expression vector of 3C protease and flipGFP were treated with rupintrivir at concentrations of 0.01, 0.03, 0.1, or 0.3 µM. The fluorescent intensity was calculated 24 h post- transfection. The data presented in E-H are representative of two independent experiments. For the experiment presented in G, significance was determined using Student’s *t*-test (n = 3) (**P ≤ 0.01; ***P ≤ 0.001; ****P ≤ 0.00001). For the experiment presented in H, significance was determined using one-way ANOVA test (n = 3) (*P ≤ 0.05; **P ≤ 0.01; ****P ≤ 0.00001; n.s., not significant).

### Characterization of polymorphism of EV-A71 3C protease using the flipGFP-based reporter system

Next, we characterized 3C proteases derived from the clinical strain of EV-A71 using the flipGFP system. The viral genome of EV-A71 has accumulated various mutations since the first isolation of the virus in 1969 in California, United States (29). We generated a phylogenetic tree of 70 unique amino acid sequences of EV-A71 3C proteases isolated from different geolocations and collection years (Fig. 5A). Additionally, the conservation index of the aligned sequence was calculated using AL2CO (30) and visualized using the reported EV-A71 structure (PDB: 8CNY) (Fig. 5B). Our conservation analysis indicated that the protein region close to the catalytic triad was highly conserved, whereas other regions sporadically changed (Fig. 5B).

**Fig. 5.**
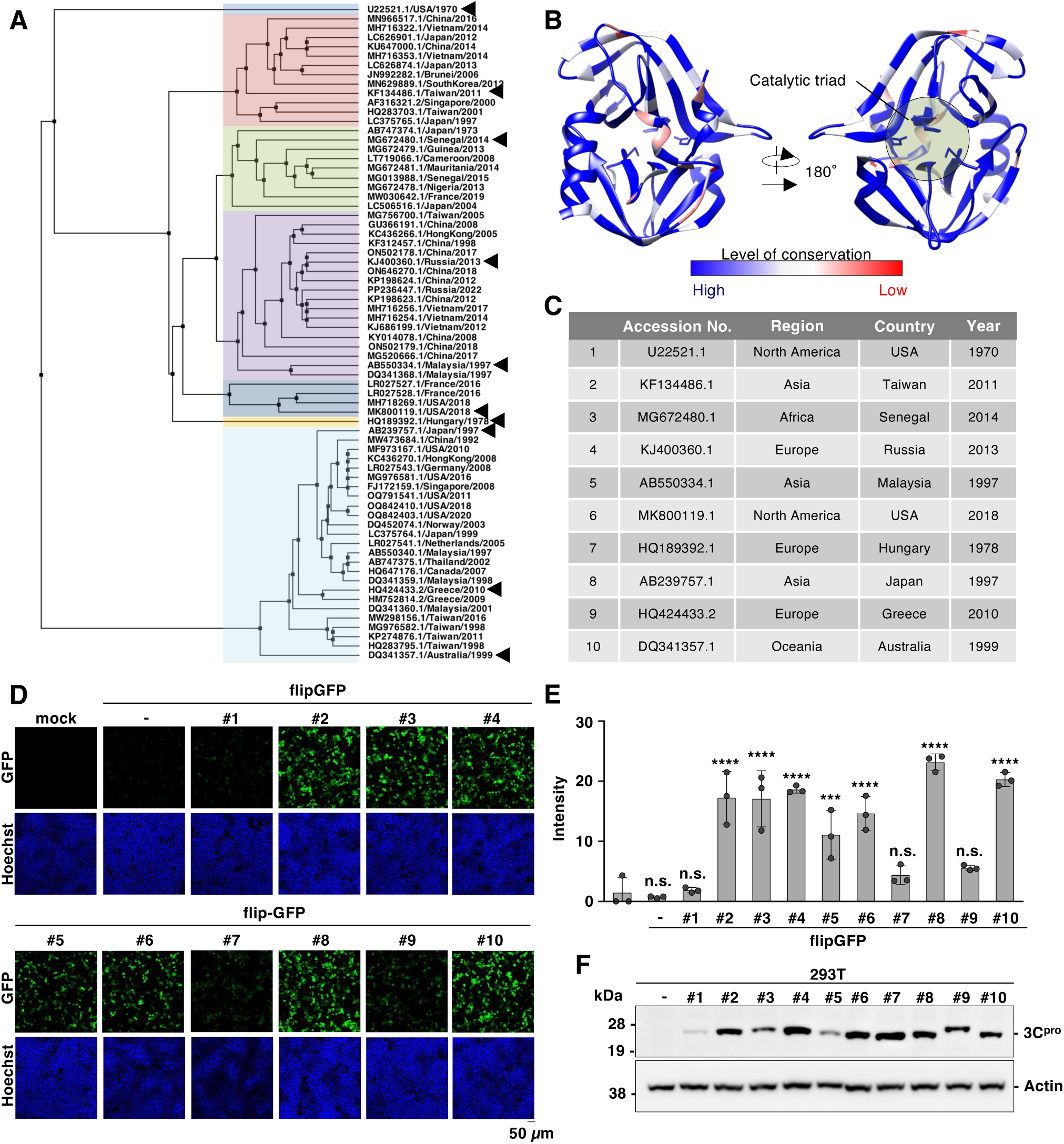
Phenotypic analysis of 3C protease derived from clinical strains of EV-A71. (A) A phylogenetic tree of 3C protease derived from clinical strains of EV-A71. The NCBI accession number, isolated country, and isolated year are indicated in each branch. The amino acid sequence of each picornaviral 3C protease was aligned by Clustal Omega (EMBL-EBI) and indicated using Jalview software version 2.11.4.1 (62). Arrowhead: the clinical strains of EV-A71 used in the subsequent study. (B) The sequences of EV-A71 3C protease presented in Fig. 5A were aligned, and the conservation level was calculated. The level of conservation was mapped in the three-dimensional structure of the 3C protease (PDB ID code: 8CNY). (C) The clinical strains of EV-A71 used in the subsequent study. (D) HEK293T cells were transfected with the plasmid encoding 3C protease derived from clinical strains of EV-A71 and the flipGFP. The fluorescent signal of flipGFP was monitored with fluorescent microscopy. (E) HEK293T cells were transfected with the plasmid encoding 3C protease derived from clinical strains of EV-A71 and the flipGFP. The fluorescent intensity was calculated 24 h post-transfection. (F) The One-strep and FLAG-tagged 3C proteases derived from clinical strains of EV- A71 were expressed in HEK293T cells. The expression of each 3C protease was detected with immunoblotting using a Strep-tag II antibody. The data presented in D, E, and F are representative of two independent experiments. For the experiment presented in E, significance was determined using one-way ANOVA (n = 3) (***P ≤ 0.001; ****P ≤ 0.00001; n.s., not significant).

These data suggest that the 3C region of the EV-A71 genome has undergone mutational changes to generate diversity during its evolution over 50 years. To further examine the phenotypic characteristics of these 3C proteases, we selected 10 clinical strains of EV-A71 that belonged to different clusters in the phylogenetic tree (Fig. 5C). First, we examined the catalytic activity of 3C proteases using the flipGFP system. The 10 amino acid sequences corresponding to the 3B/3C region were identical among the strains (Fig. S3A). We co-expressed flip-GFP with each 3C protease in HEK293T cells. A significant fluorescent signal was observed in seven strains of EV-A71 examined, whereas a reduced signal was observed in strains #1, #7, and #9 (Fig. 5D and E). Immunoblotting analysis of the 3C proteases derived from strains #7 and #9 confirmed that their expression was comparable to that of other strains, such as #10, while strain #1 showed weaker expression (Fig. 5F). We focused on strains #7 and #9 and sought to identify the amino acid residues responsible for 3C protease activity.

The aligned sequences of the 3C proteases showed 11 differences in amino acid residues between strains #7 and #10, and five between strains #9 and #10 (Fig. S3B). Therefore, we generated a series of mutant 3C proteases that swapped the amino acid residues of strains #10–#7 (H33R, K55N, V57L, T93I, P96G, G145W, A153S, V157I, A173G, G177S, and C180A) and #9 (Q24H, I68V, I86T, K88S, and N111S) (Fig. 6A). We co-expressed the mutant 3C proteases with flipGFP in HEK293T cells and observed a significant decrease in the signal in the G145W or I86T mutants (Fig. 5B-E). Conversely, substitution of W145 in the 3C protease (strain #7) with G (Fig. 5F) or T86 in the 3C protease (strain #9) with I (Fig. 5G) recovered the flipGFP signal (Fig. 5H, I, J, and K). These results suggest that the two specific amino acid residues located at residues 86^th^ and 145^th^ of the 3C protease are crucial for its protease activity. The three-dimensional structure of the EV-A71 3C protease indicates that the 145^th^ amino acid position is located near the catalytic triad, whereas the 86^th^ amino acid position is not (Fig. 6L), suggesting that the significance of these two amino acids is distinct.

**Fig. 6.**
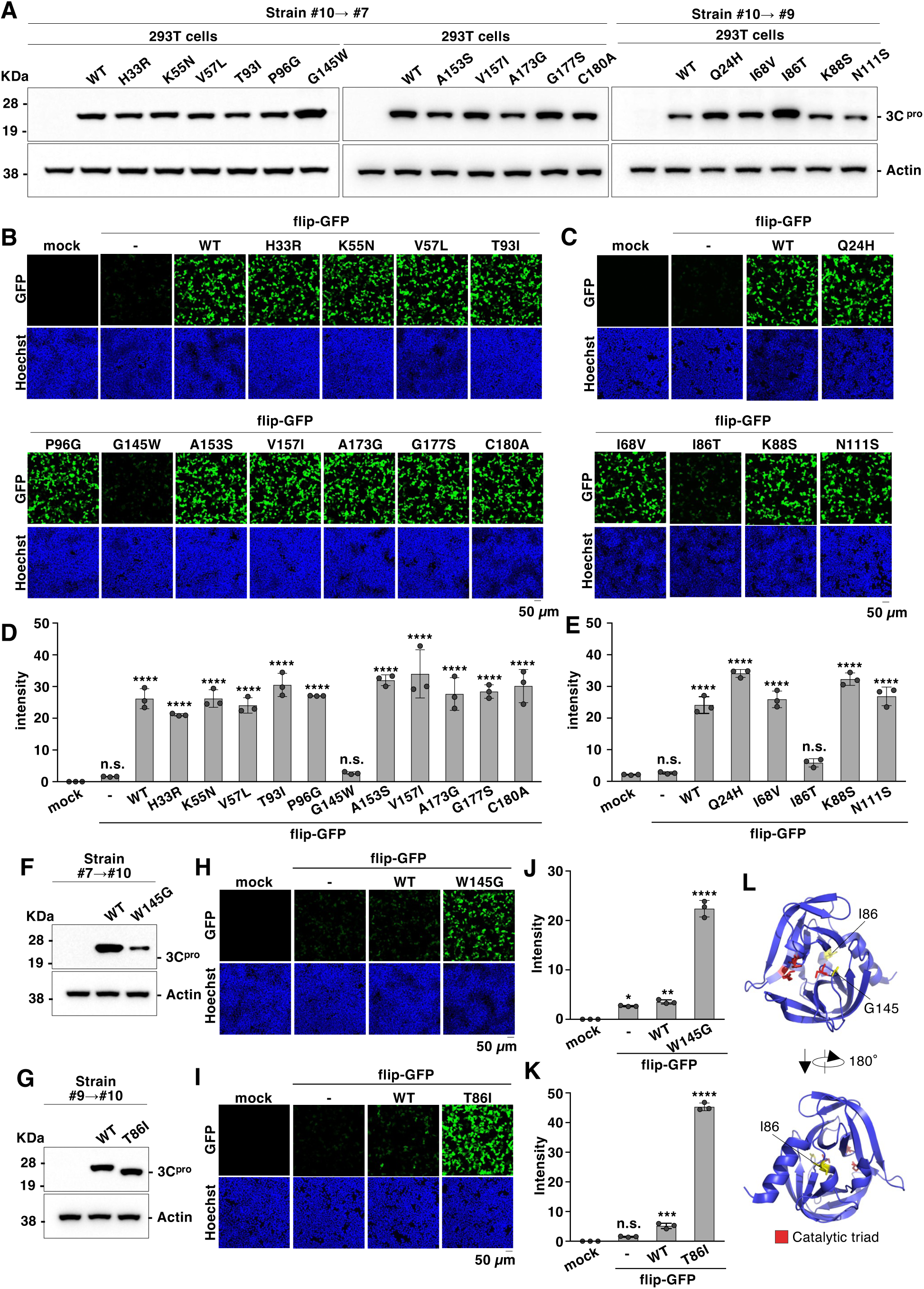
Eighty-sixth and 145^th^ amino acids in the EV-A71 3C protease influence its activity. (A) Each amino acid residue of clinical strain EV-A71 #10, which is distinct from #7 or #9, was mutated into the corresponding amino acid. Mutated 3C proteases were expressed in HEK293T cells and detected with immunoblotting using a Strep-tag II antibody. (B-C) HEK293T cells were transfected with a plasmid encoding the mutated 3C protease and flipGFP. The fluorescence signal of flipGFP was monitored using fluorescence microscopy. (D-E) HEK293T cells were transfected with plasmids encoding the mutated 3C protease and flipGFP. The fluorescence intensity was calculated 24 h post-transfection. (F) The mutated 3C protease carrying a tryptophan to glycine substitution at the 145^th^ amino acid position (W145G) was expressed in HEK293T cells and detected by immunoblotting using a Strep-tag II antibody. (G) The mutated 3C protease, carrying a substitution from threonine to isoleucine at the 86^th^ amino acid position (T86I), was expressed in HEK293T cells and detected with immunoblotting using a Strep-tag II antibody. (H) HEK293T cells were transfected with plasmids encoding a mutated 3C protease (W145G) and flip-GFP. The fluorescence signal of flipGFP was monitored using fluorescence microscopy. (I) HEK293T cells were transfected with plasmids encoding a mutated 3C protease (T86I ) and flip-GFP. The fluorescence signal of flipGFP was monitored using fluorescence microscopy. (J) HEK293T cells were transfected with plasmids encoding a mutated 3C protease (W145G) and flip-GFP. The fluorescence intensity was calculated 24 h post-transfection. (K) HEK293T cells were transfected with plasmids encoding a mutated 3C protease (T86I) and flip-GFP. The fluorescence intensity was calculated 24 h post-transfection. (L) The amino acid residues located at the 86^th^ and 145^th^ positions in the 3C protease are yellow in the three-dimensional structure of the 3C protease (PDB ID code: 8CNY). Data presented in A-K are representative of two independent experiments. For the experiments presented in D, E, J, and K, significance was determined using one- way ANOVA (n = 3) (*P < 0.05; **P < 0.01; ****P < 0.00001; n.s., not significant).

### The cleavage activity of host proteins by 3C protease differed among EV-A71 clinical strains

To further determine how the genetic polymorphism of the 3C protease influences the actual EV-A71 characteristics, we generated a series of chimeric EV-A71 strains in which the 3C protease region was swapped with other strains of EV-A71 (Fig. 7A). Using strain SK-EV006 (strain #5) (14) as the backbone, infectious chimeras were recovered from strains that confirmed the catalytic activity of the 3C protease using flipGFP (Fig. 5D and E). The generated passage 0 (P0) viruses showed comparable titers and plaque phenotypes (Fig. 7B and C). Conversely, the infectious virus was not recovered from strains #1, #7, or #9 (Fig. 7B). As the 3C protease cleaves multiple host proteins, including G3BP1 and eIF5B (19, 20), we sought to determine the cleavage activity of host proteins by these chimeras. The cleaved form of G3BP1 or eIF5b was observed after infection with these chimeras, whereas the amount of cleaved products differed among the strains (Fig. 7D). To validate these results, we focused on chimeras carrying the 3C protease derived from strains #8 and #10 (SK-#8 and SK-#10, respectively).

**Fig. 7.**
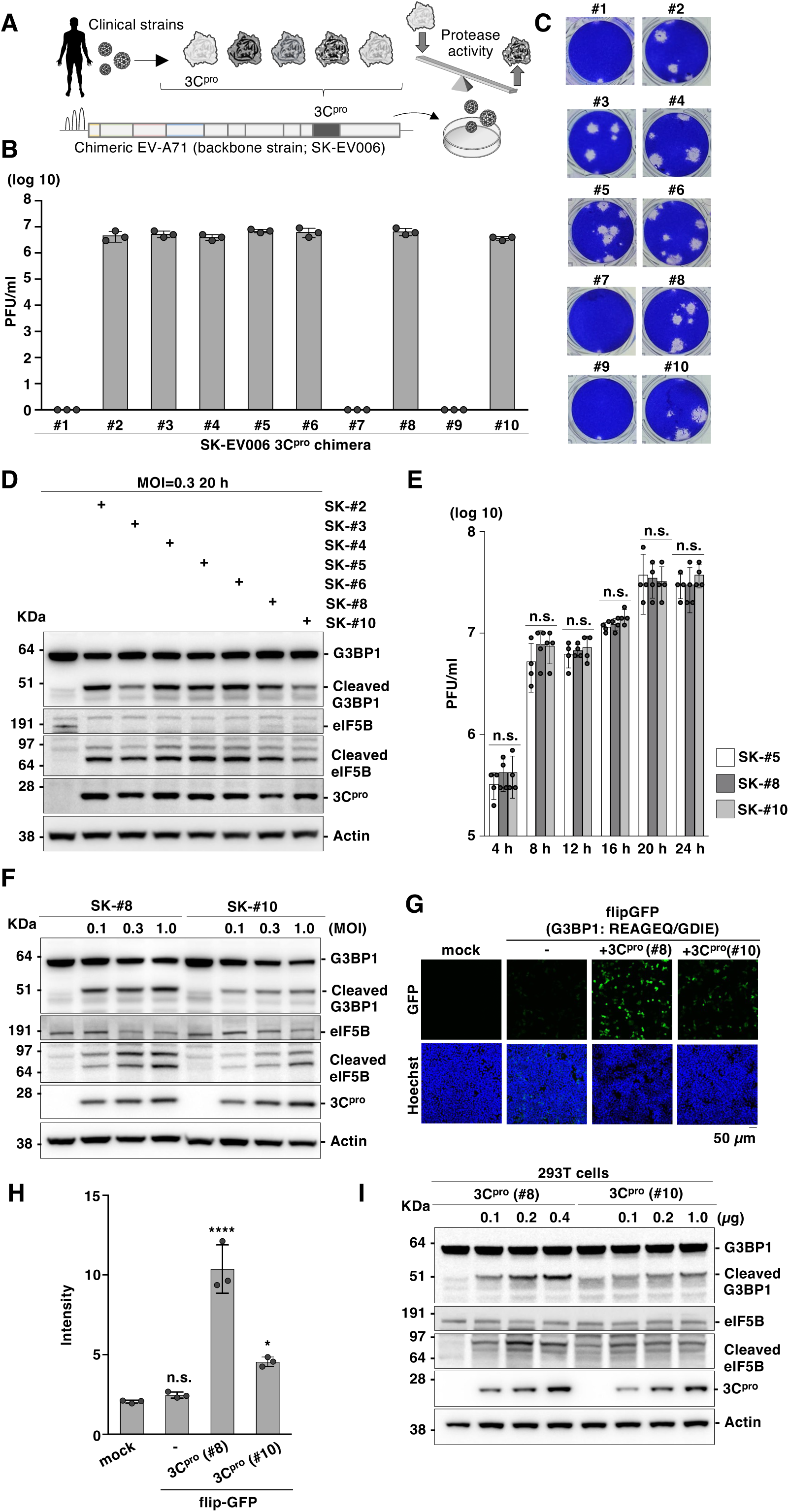
Clinical strains of EV-A71 cleave host proteins differently. (A) Schematic representation of the chimeric EV-A71 carrying a clinical strain of 3C protease. The 3C protease coding region in the EV-A71 genome (strain SK-EV-006) was swapped to the sequence derived from the clinical strain of 3C protease (strains #1–#10). The chimeric EV-A71 was generated by a reverse genetics system for EV-A71. (B) Infectious titers of the generated chimeric EV-A71 were determined using a plaque-forming assay. (C) Plaque morphology of chimeric EV-A71. The dilution of the viruses was 10^−6^. (D) Expression of G3BP1 and eIF5b in RD-S cells infected with chimeric EV- A71 at an MOI of 0.3 was detected using immunoblotting at 20 h post-infection. (E) RD-S cells were infected with EV-A71 carrying 3C protease of wild-type (SK-#5), strain #8 (SK-#8) or strain #10(SK- #10) at an MOI of 1.0, and infectious titers were determined using a plaque-forming assay at 4, 8, 12, 16, 20, or 24 h post-infection. (F) Expression of G3BP1 and eIF5b in RD-S cells infected with chimeric EV-A71 at an MOI of 0.1, 0.3, or 1.0 was detected using immunoblotting at 20 h post-infection. (G) HEK293T cells were transfected with the plasmid encoding 3C protease of strain#8 (3C^pro^ (#8)) or #10 (3C^pro^ (#10)) and the flipGFP inserted sequence derived from G3BP1. The fluorescent signals were detected using microscopy at 24 h post-transfection. (H) HEK293T cells were transfected with the plasmid encoding the 3C protease of strain #8 or #10 and the flipGFP inserted sequence derived from G3BP1. The fluorescent intensity was calculated 24 h post-transfection. (I) HEK293T cells were transfected with increasing amounts of a plasmid encoding the 3C protease of strain #8 or #10, and G3BP1 and eIF5b expression were detected using immunoblotting at 24 h post-transfection. Data presented in E as mean ± S.D. of two independent experiments; data in B-D and F-I are representative of two independent experiments. For the experiment presented in E and H, significance was determined using one-way ANOVA test (n = 3 or 4) (*P ≤ 0.05; ****P ≤ 0.00001; n.s., not significant).

First, a one-step growth analysis was performed, confirming that these two chimeras propagated equally to the wild-type EV-A71 SK-EV006 (strain #5) (Fig. 7E).

Next, the cells were infected with different multiplicities of infection (MOI) of SK-# 8 or SK-# 10 to observe cleaved G3BP1 and eIF5b. The cleaved products were readily observed in SK-#8 infected cells and moderately in SK-#10 infected cells, even at a high MOI (Fig. 7F). To comprehensively evaluate the effect of different cleaving activities of the protease during EV-A71 infection, RNA sequencing (RNA-seq) was performed in cells infected with the chimeric virus carrying strain #8 or #10 of the 3C protease (Fig. S4A). Comparison of the RNA-seq profiles between the cells infected with the wild-type (SK-#5) and chimeric (SK-#8 or SK-#10) viruses showed an overall similarity in gene expression patterns; however, differences, such as the pathway related to the morphology of the nucleus, were observed among the compared profiles between SK-#8 and SK-#10 (Fig. S4B). These data suggest that the cellular responses to these viruses differ. To functionally validate and quantitatively determine whether the 3C protease alone was responsible for the observed phenotype, we constructed a flipGFP carrying the G3BP1 sequence targeted by the 3C protease (REAGEQ/GDIE, flipGFP-G3BP1) (20).

We co-expressed flipGFP-G3BP1 with 3C protease (strains #8 or #10) and observed a significant signal, especially in cells co-expressing both flipGFP-G3BP1 and strain #8 3C protease (Fig. 7H). Conversely, a comparable fluorescent signal was observed when strain #8 or #10 of the 3C protease was co-expressed with flipGFPs encoding the viral cleavage sequence (VP2/VP3, VP3/VP1, 2C/3A, or 3C/3D) (Fig. S3C and D). We also constructed flipGFP encoding the eIF5B sequence (VEVMEQ/GVPE) targeted by the 3C protease; however, detection of the signal was hampered by the high background signal. Therefore, we expressed 3C protease (strain #8 or #10) in HEK293T cells and confirmed the cleavage activity of the endogenous substrate using immunoblotting. Consistent with the results presented in Figure 7 F, endogenous G3BP1 or eIF5B was preferentially cleaved during the expression of the 3C protease in strain #8, reproducing the results of the chimeric viruses (Fig. 7I).

Taken together, our data combining flipGFP and the chimeric viruses suggest that the ability of the 3C protease to cleave viral protein and produce infectious particles is mostly conserved, while the ability to cleave host protein that affect cellular responses can vary among the strains.

## Discussion

Cell-based reporters applicable to a large number of genetically heterogeneous viral families, such as Picornaviridae, remain lacking. This study addresses this knowledge gap by developing a novel flipGFP-based protease assay for human pathogenic picornaviruses (genera *Enterovirus*, *Rhinovirus*, *Cosavirus*, *Salivirus*, and *Kobuvirus*), as well as previously neglected picornaviruses derived from viral reservoirs (bat, simian, and rodent picornaviruses). Subsequently, we determined how the polymorphism of EV-A71 3C protease influenced its cleavage activity using a flipGFP reporter. By comparing the clinical strains of EV-A71, we found that amino acid residues influenced the 3C protease activity. Furthermore, using flipGFP and chimeric EV-A71 expressing different 3C proteases, our findings demonstrated that polymorphisms in the EV-A71 3C protease influenced the cleavage activity of host proteins. Our experiments demonstrate how flipGFP enhances our understanding of the molecular characteristics of picornaviral 3C proteases.

Among the eight sequences within the EV-A71 polyprotein targeted by the 3C protease, the sequence between VP2/VP3 or 3B/3C was optimal for the flipGFP assay. In contrast, flipGFP encodes a sequence derived from a relatively stable precursor protein, especially the sequence between 2B/2C or 3A/3B, barely emitting the fluorescent signal in the presence of 3C protease. Furthermore, we showed that the sequence between 3B/3C could be used to develop flipGFP for a wide range of picornaviruses, suggesting that the efficient cleavage of this sequence is conserved among Picornaviridae. Therefore, this sequence could be used to develop flipGFP for the detection of unknown picornaviruses in the future. Conversely, inserting the sequence between VP2 and VP3 into flipGFP is viable for detecting the activity of 3C proteases derived from EV-A71 or EV-D68 but not from CVB3 or PV1. The observed results suggest that the efficacy of cleaving structural regions by the 3C protease, which balances particle production and viral replication, differs among enterovirus species. Further studies are warranted to precisely determine how this difference influences the characteristics of the viruses and the associated biological significance.

Since our flipGFP reporter reflects the inhibitory activity of a known 3C protease inhibitor, drug screening could be applicable to picornaviral 3C proteases. Consistently, at the time of the COVID-19 pandemic, the flipGFP reporter was rapidly optimized to detect the activity of the main protease (M^pro^) derived from many coronaviruses, including the SARS-CoV-2 (31). The reporter of M^pro^ was subsequently used for high-throughput drug screening to identify inhibitors against SARS-CoV-2.

Several novel M^pro^ inhibitors have been reported, including those by our group (32–37). Compared with the conventional *in vitro* recombinant assay, the flipGFP protease assay can flexibly change the enzyme and reflect the actual cellular environment, where the protease is located in complex heterogeneity and co-exists with several co-factors. Inhibitors of previously uncharacterized 3C proteases, such as animal picornaviruses, can provide novel insights into existing 3C protease inhibitors and should be further explored.

To demonstrate the flexibility of flipGFP in comparing protease activity within viral species, we examined the differences in the 3C proteases of clinical EV-A71 strains. Using flipGFP, we showed that the sequence between 3B and 3C is highly conserved among viruses belonging to different clusters in the phylogenetic tree of 3C proteases, and the sequence is cleaved comparably by 7 out of the 10 proteases we examined. Conversely, some proteases exhibit low protease activity regulated by specific amino acid residues, such as the 86^th^ and 145^th^ amino acid positions in the 3C protease. Structural analysis of the protease predicted that the substitution of glycine with threonine at the 145^th^ amino acid position in the 3C protease reduced access of the substrate to the catalytic triad, suggesting that this mutation directly influences catalytic activity. In contrast, the 86^th^ amino acid was located diametrically opposite to the catalytic triad in the structure. This region binds viral RNA, supporting viral replication (38–40). Therefore, the polymorphism of the 86^th^ amino acid in the 3C protease might regulate the functional balance between cleavage and replication of the 3C protease. Although we could not confirm the propagation of chimeric EV-A71 strains carrying T86 or W145 (Fig. 7B), these mutations could be selected and made viable in a specific *in vivo* environment, the significance of which should be further characterized using authentic clinical strains.

The chimeric EV-A71 strains carrying different strains of 3C protease showed comparable titers and plaque phenotypes, suggesting that the function of the 3C protease in cleaving viral polyproteins to support viral propagation is conserved. In contrast, we observed differences in the activity of proteases that cleave host proteins. Specifically, the 3C protease detected in the HFMD case in Japan (strain #8) (41, 42) exhibited high cleavage activity toward the host G3BP1 or eIF5b protein, whereas the protease from the meningitis case in Australia (#10) (43) barely cleaved these proteins. The relationship between the cleavage activity of host proteins by the 3C protease and the induction of EV-A71 pathogenesis remains unclear; this is the first report to demonstrate that cleavage events by the protease differ among EV-A71 strains. G3BP1 is involved in the formation of stress granules and acts as an antiviral protein during infection (44). Although enterovirus infection inhibits stress granule formation, its function is mainly mediated by the 2A protease (45, 46). Therefore, G3BP1 cleavage by the 3C protease may regulate the noncanonical function of G3BP1 to support viral propagation (47). Cleavage of eIF5b by 3C protease contributes to the shutoff of translation, thereby evading host antiviral responses (19). The 3C protease cleaves a large number of host proteins, including G3BP1 and eIF5b, to propagate efficiently (48); however, the mechanism by which these cleavages influence the host or virus remains nebulous. In this study, we inserted the host-derived G3BP1 sequence into the flipGFP and confirmed that the signal depended on protease activity. Since flipGFP is highly flexible in changing the cleavage sequence inside the protein, constructing a flipGFP library covering a wide range of host substrates will expand our understanding of the cleavage patterns of host proteins in various enterovirus strains and aid in conducting comparisons.

Although our assay is a robust tool for evaluating the characteristics of 3C proteases, some limitations remain in the current system. Specifically, a high background signal inhibited the flip-GFP assay for eIF5G, suggesting that the sequence is cleaved by cellular proteases. Therefore, further optimization of flipGFP is required to thoroughly evaluate host cleavage. The flipGFP reporter for SARS-CoV-2 was later improved to reduce the background signal by adding the FKBP12 degron (49). This modification may enhance the picornaviral flipGFP reporter. In addition, our system did not normalize the expression level of the 3C protease, which showed some variation and could have influenced the results. Therefore, combining multiple assays to examine the results is warranted.

In this study, we present a novel cell-based reporter adjustable for diverse picornaviral 3C proteases. The reporter allows for the screening of therapeutics for well-characterized and previously neglected picornaviruses. Additionally, our report can aid in developing preventive measures against future picornaviruses. This study also demonstrated the utility of this reporter for evaluating genetic polymorphisms of proteases that influence the cleaving activity of host proteins. Therefore, our reporter could prove useful for future comprehensive studies aimed at understanding the pathogenic mechanisms of picornaviruses associated with the 3C protease activity.

## Materials and methods

### Cell lines and viruses

Human embryonic kidney (HEK293T) and rhabdomyosarcoma (RD) cell lines were obtained from the National Institute of Infectious Diseases, Japan Institute for Health Security, Tokyo, Japan. The cell lines were cultured in Dulbecco’s Modified Eagle Medium (DMEM; High Glucose) with L-Glutamine and Phenol Red (Wako, 044-29765) supplemented with 10 % fetal bovine serum (FBS), 100 U/mL penicillin, and 100 µg/mL streptomycin. EV-A71 (strain SK-EV006) (14) and chimeric EV-A71 carrying a 3C protease from clinical strains were generated using a reverse genetics system for EV- A71, as previously described (50, 51). Briefly, the pcDNA3 vector encoding the EV-A71 genome was transfected into RD cells using the TransIT-LT1 Transfection Reagent (Mirus) according to the manufacturer’s protocol. The culture supernatant and cells were collected at 3 days post-transfection, freeze-thawed three times, and centrifuged at 13,000 rpm for 5 min at 4 ℃. The supernatants were collected, kept at –80 ℃, and used for the study.

### Antibodies and reagents

The following antibodies and reagents were used in this study: Enterovirus 71 3C antibody (GTX132357; GeneTex), anti-FLAG M2-Peroxidase (HRP) monoclonal antibody produced in mouse (A8592; Merck), Anti-Strep-tag II mAb (M211-3; MBL), Anti-DDDDK-tag mAb-Alexa Fluor™ 594 (M185-A59, MBL), anti-β-Actin monoclonal antibody produced in mouse (A2228; Merck), Peroxidase AffiniPure Goat anti-mouse IgG (H+L) (115-035-003; Jackson Immuno Research Laboratories, Inc.), Peroxidase AffiniPure Goat anti-Rabbit IgG (H+L) (111-035-003; Jackson Immuno Research Laboratories, Inc.), and Rupintrivir (T16809; TargetMol).

### Plasmids

PCDNA3-FlipGFP(Casp3 cleavage seq) T2A mCherry was a gift from Xiaokun Shu (Addgene plasmid # 124428; http://n2t.net/addgene:124428; RRID:Addgene_124428). pUltra was gifted by Malcolm Moore (Addgene plasmid # 24129; http://n2t.net/addgene:24129; RRID:Addgene_24129). pMDLg/pRRE and pRSV-Rev were gifts from Didier Trono (Addgene plasmids # 12251 and # 12253; http://n2t.net/addgene:12251 and http://n2t.net/addgene:12253; RRID:Addgene_12251 and RRID:Addgene_12253, respectively). pCMV-VSV-G was a gift from Bob-Weinberg (Addgene plasmid # 8454; http://n2t.net/addgene:8454; RRID:Addgene_8454). The cDNA of flipGFP was amplified using PCR, cloned into the pCAGGS vector (52) digested with the EcoRI suite. Sequences of the protease cleavage sites inside flipGFP are indicated in each figure. The synthesized cDNA of picornaviral 3C proteases was amplified using PCR by adding N-terminal One-Strep and FLAG tags. The sequence information of each picornaviral protease is indicated in each figure as a GenBank accession number. The 3C protease sequence derived from CV-A6, Echovirus, HRV-A, and Aichhivirus has been codon optimized in humans to enhance its expression. The amplified cDNA of picornaviral 3C proteases was cloned into the pCAGGS vector and digested using the EcoRI suite. All cloning procedures were conducted using an In-Fusion HD cloning kit (Clontech) or In-Fusion Snap Assembly Master Mix (Clontech). Sequences of the generated plasmids were confirmed using a SeqStudio Genetic Analyzer (Thermo Fisher Scientific).

### flipGFP assay

HEK293T cells (1.5×10^5^ cells in 0.25 mL of culture medium) seeded on a 24-well plate were incubated at 37 °C for 24 h and transfected with 0.25 µg of pCAGGS-flipGFP vector and 0.025 µg of pCAGGS EV-A71 3C protease vector using TransIT-LT1 Transfection Reagent (Mirus) according to the manufacturer’s protocol. The transfected cells were incubated for 16 h and inoculated with an additional culture medium (0.25 mL) containing Hoechst 33342 (H1399; Thermo Fisher Scientific) at a concentration of 10 µg/mL. The fluorescent signals of flipGFP were observed using a microscope (FV- 3000; OLYMPUS) and quantified using CellProfiler software (Cimini Lab, the Broad Institute of MIT and Harvard).

### Immunoblotting

Cells were lysed in a lysis buffer consisting of 20 mM Tris⋅HCl (pH 7.4), 135 mM NaCl, 1 % Triton X- 100, 1 % glycerol, and protease inhibitor mixture tablets (Roche), incubated on ice for 30 min, and centrifuged at 15,000 rpm for 5 min at 4 ℃. The supernatants were collected from the cell lysate and supplemented with an equal volume of sodium dodecyl sulfate (SDS) gel-loading buffer (2×) containing 50 mM Tris-HCl (pH = 6.8), 4 % SDS, 0.2 % bromophenol blue, 10 % glycerol, and 200 mM β-mercaptoethanol. The mixture was subsequently boiled at 95 °C for 5 min, resolved using SDS polyacrylamide gel electrophoresis (NuPAGE gel, Thermo Fisher Scientific), and transferred onto nitrocellulose membranes (iBlot2, Thermo Fisher Scientific). The membranes were incubated with primary antibodies diluted in PBS containing 0.05 % Tween20 (PBST) supplemented with 5 % skim milk (1:1,000 dilution) at 4 °C for 24 h. The membranes were then washed thrice with PBST, incubated with secondary antibodies (1:3,000 dilution) at room temperature for 1 h, and washed thrice with PBST. The blots were visualized using Amersham ECL western blotting detection reagent (GE Healthcare) and an Amersham Imager (GE Healthcare).

### Immunofluorescence staining

HEK293T cells (6.0×10^4^ cells in 0.3 mL of culture medium) were seeded on a cover glass chamber (IWAKI, 5232-008), incubated at 37 ℃ for 24 h, and transfected with the plasmid encoding picornaviral 3C protease using TransIT-LT1 Transfection Reagent (Mirus) according to the manufacturer’s protocol. The cells were incubated for 24 h, fixed with 4 % paraformaldehyde in PBS at room temperature for 30 min, and incubated with 0.2 % Triton X-100 in PBS for 25 min. The cells were subsequently incubated with 5 % FBS in PBS for 1 h and incubated with Anti-DDDDK-tag mAb-Alexa Fluor™ 594 (M185-A59, MBL) (1:1,000 dilution) at 4 ℃ for 24 h. Subsequently, the cells were incubated with Cellstain DAPI solution (Dojindo, D523) (1:5,000 dilution) at room temperature for 15 min, and the fluorescence was analyzed using microscopy (FV-3000; OLYMPUS) and cellSens software (OLYMPUS).

### Establishment of RD cells stably expressing SCARB2

For the experiments in Figure 7B-F, RD cells lentivirally expressing SCARB2 (53, 54) were generated. Plasmid pUltra (Addgene plasmid # 24129) was used as the backbone of the lentiviral transfer vector. The puromycin N-acetyltransferase sequence under the encephalomyocarditis virus internal ribosomal entry site was inserted into pUltra, which was digested with AgeI and SalI. The resulting plasmid, RUIPW, was subsequently digested with BamHI, and the SCARB2 sequence synthesized by Eurofins Japan was inserted. The resultant plasmids (RUIPW SCARB2), pMDLg/pRRE (Addgene plasmid # 12251), pRSV-Rev (Addgene plasmid #12253), and pCMV-VSV-G (Addgene plasmid # 8454) were transfected into HEK293T cells using TransIT-LT1 Transfection Reagent (Mirus) according to the manufacturer’s protocol. The culture supernatant containing lentivirus was collected 3 days post-transfection and passed through a 0.45-µm filter. For the infection of lentivirus, the seeded RD cells were inoculated with the generated lentivirus supplemented with hexadimethrine bromide (4 μg/mL; Sigma). Followed by the centrifugation at 2,500 rpm for 50 min at 32 °C, the cells were cultured for 3 days and selected by the addition of puromycin at a concentration of 1 µg/mL. Selected RD cells expressing SCARB2 were named RD-S cells.

### Plaque formation assay

RD-S cells (7.5 × 10^4^ cells in 0.25 µL of culture medium) seeded on a 48-well plate were incubated at 37 ℃ for 24 h. The cells were inoculated with the serially diluted EV-A71, washed three times with DMEM supplemented with 2 % FBS, and incubated in DMEM supplemented with 0.3 % cellulose (MERCK, 435244) and 10 % FBS for 3 days. The Plaques were visualized using 20 % ethanol in PBS containing 0.5 % crystal violet.

### Analysis of the structure and recognition sequence of 3C protease

For the experiment presented in Figures 4C and D, WebLogo Version 2.8.2 (https://weblogo.berkeley.edu/logo.cgi) was used to analyze the sequence conservation of amino acids (55, 56). For the experiment presented in Figure 5B, the conservation level of 3C protease was visualized as a three-dimensional structure using UCSF ChimeraX version 1.9, developed by the Resource for Biocomputing, Visualization, and Informatics at the University of California, San Francisco, with support from the National Institutes of Health R01-GM129325 and the Office of Cyber Infrastructure and Computational Biology, National Institute of Allergy and Infectious Disease (57–59). For the experiment presented in Figure 6L, the three-dimensional structure of the 3C protease was visualized using PyMOL (open-source PyMOL Molecular Graphics System 1.8.4.0).

### RNA sequencing (RNA-seq)

RD-S cells were infected with chimeric EV-A71 at an MOI of 1.0, incubated for 16 h, and total RNA was extracted using ISOGEN II (Nippon Gene) according to the manufacturer’s protocol. The TruSeq stranded mRNA sample prep kit (Illumina, San Diego, CA, USA) was used to prepare the sequencing libraries. The resulting library was sequenced using an Illumina NovaSeq 6000 platform in 100-bp single-end mode. The acquired sequence reads were mapped to human reference genome sequences (hg19) using TopHat v2.0.13, in combination with SAMtools ver. 0.1.19. Cufflinks version 2.2.1 and Bowtie2 ver. 2.2.3. To calculate the fragments per kilobase of exons per million mapped fragments (FPKMs), Cufflinks version 2.2.1. was used. RNA-seq data were visualized using the iDEP 2.01 server (http://bioinformatics.sdstate.edu/idep/) (60). GSEA (preranked fgsea) method and Gene Ontology (GO) Molecular Function Pathway database were used to analyze the biological pathway presented in Figure 4B.

### Statistical analysis

Statistical analyses presented in the figures were conducted using GraphPad Prism version 9.5.1 (GraphPad). Significant differences were calculated using Student’s *t*-test or one-way analysis of variance (ANOVA) test. The details of the analyses employed in each experiment are described in the figure legends.

## Acknowledgements

We thank M. Tomiyama, T. Toyoshima, M. Ishibashi, and K. Toyoda for their assistance. We also thank all the members of the Department of Virology II, National Institute of Infectious Diseases, Japan Institute for Health Security, and the Laboratory of Virus Control, University of Osaka, for their kind support. This work was partly supported by the Ministry of Education, Culture, Sports, Science, and Technology of Japan (KAKENHI) (grant numbers 22K15479, 22K06625, 24K02286, and 25K18815), Takeda Science Foundation (grant number 2024047859), the Japan Agency for Medical Research and Development (AMED) (grant number JP25fk0108729), and the Japan Science and Technology Agency (JST) (Moonshot R&D) (JPMJMS2025). This research was conducted as part of the All-Osaka U Research in "The Nippon Foundation - Osaka University Project for Infectious Disease Prevention.”

## Data Availability

All data are shown in the main figures and supplementary material. Materials, microscopic images, sequence information, and raw sequencing data are available upon request. Requests for materials and correspondence should be addressed to Professor Yoshiharu Matsuura.

## Author Contributions

Conceptualization, J.H., Y.S., and Y.M.; Methodology, J.H. and Y.M.; Investigation, J.H., T.H., Y.S., K.O., K.U., M.Y., C.O., and S.T.; Resources, J.H., T.H., Y.S., and Y.M.; Writing-Original Draft, J.H. and Y.M.; Writing-Review and Editing, T.H., Y.S., and Y.M.; Funding Acquisition, J.H., T.H., Y.S., and Y.M.; Supervision, Y.M.

---

**Supplementary Figure 1.**
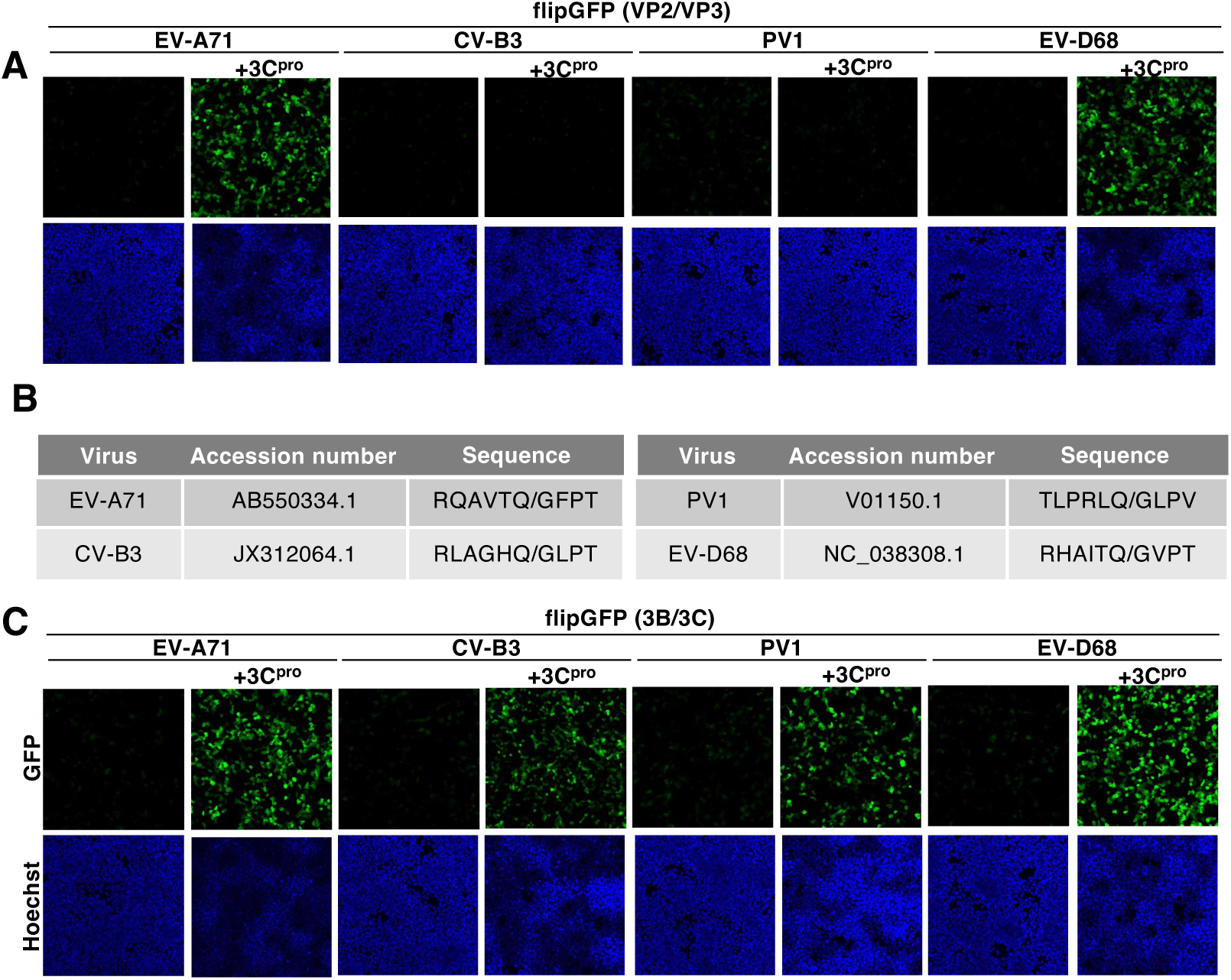
(A) HEK293T cells were transfected with the plasmid encoding 3C protease derived from genus *Enterovirus* (EV-A71, CVB3, PV1, and EV-D68) and the flipGFP (VP2/VP3). The fluorescent signal of flipGFP was monitored with fluorescent microscopy. (B) The accession number of the *Enterovirus* used in this study and the corresponding amino acid sequence between the VP2 and VP3 regions. (C) HEK293T cells were transfected with the plasmid encoding 3C protease derived from the genus *Enterovirus* (EV-A71, CVB3, PV1, and EV-D68) and the flipGFP (3B/3C). The fluorescent signal of flipGFP was monitored with fluorescent microscopy. The data presented in A and C are representative of two independent experiments.

**Supplementary Figure 2.**
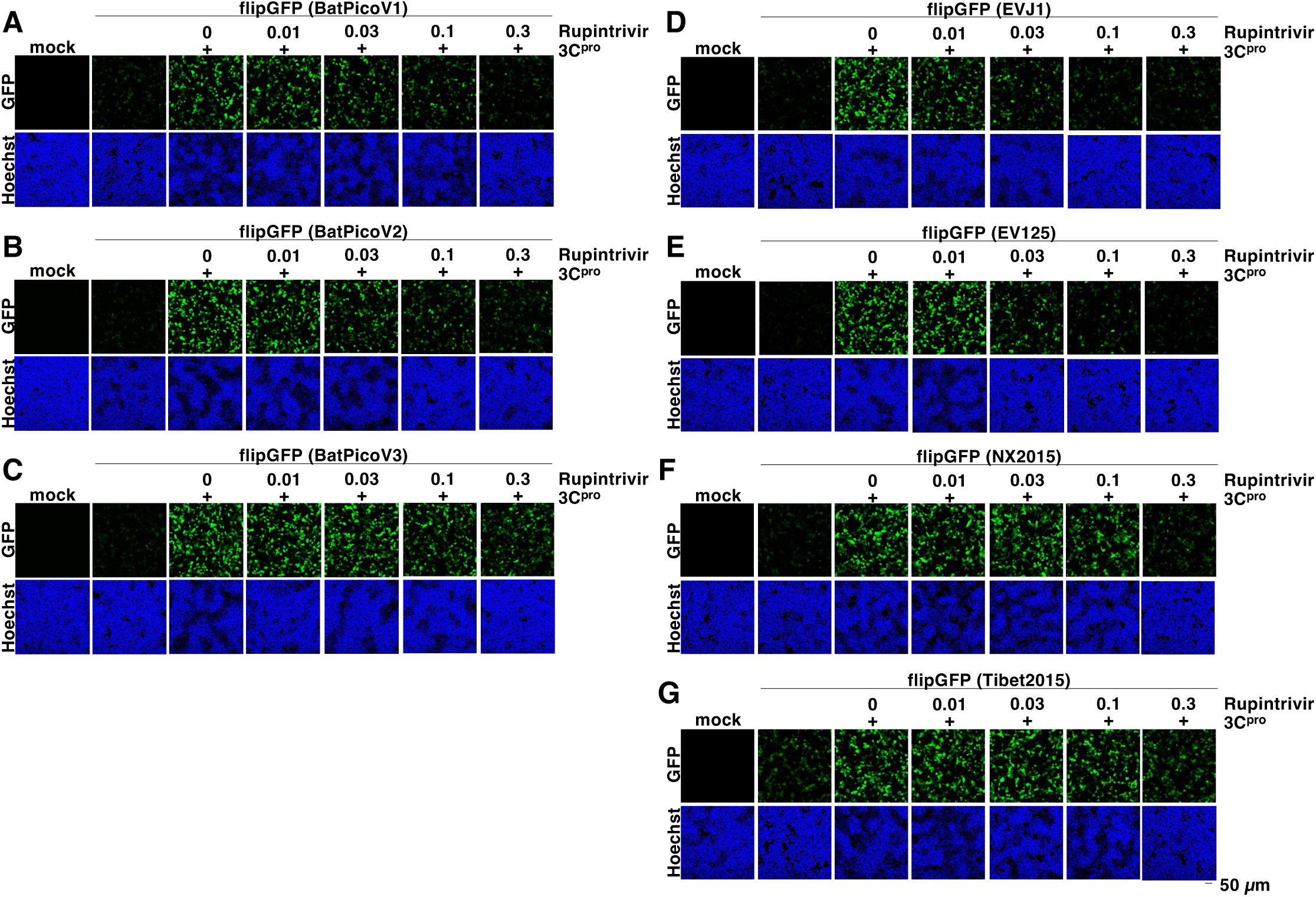
(A-H) HEK293T cells transfected with the expression vector of 3C protease and flipGFP (BatPicoV1 for Fig. S2A, BatPicoV2 for S2B, BatPicoV3 for Fig. S2C, EVJ1 for Fig. S2D, EV125 for Fig. S2E, NX2015 for Fig. S2F, and Tibet2015 for Fig. S2G) were treated with rupintrivir at concentrations of 0.01, 0.03, 0.1, or 0.3 µM. The fluorescent signals were detected using microscopy at 24 h post-transfection. The data presented in A-H are representative of two independent experiments.

**Supplementary Figure 3.**
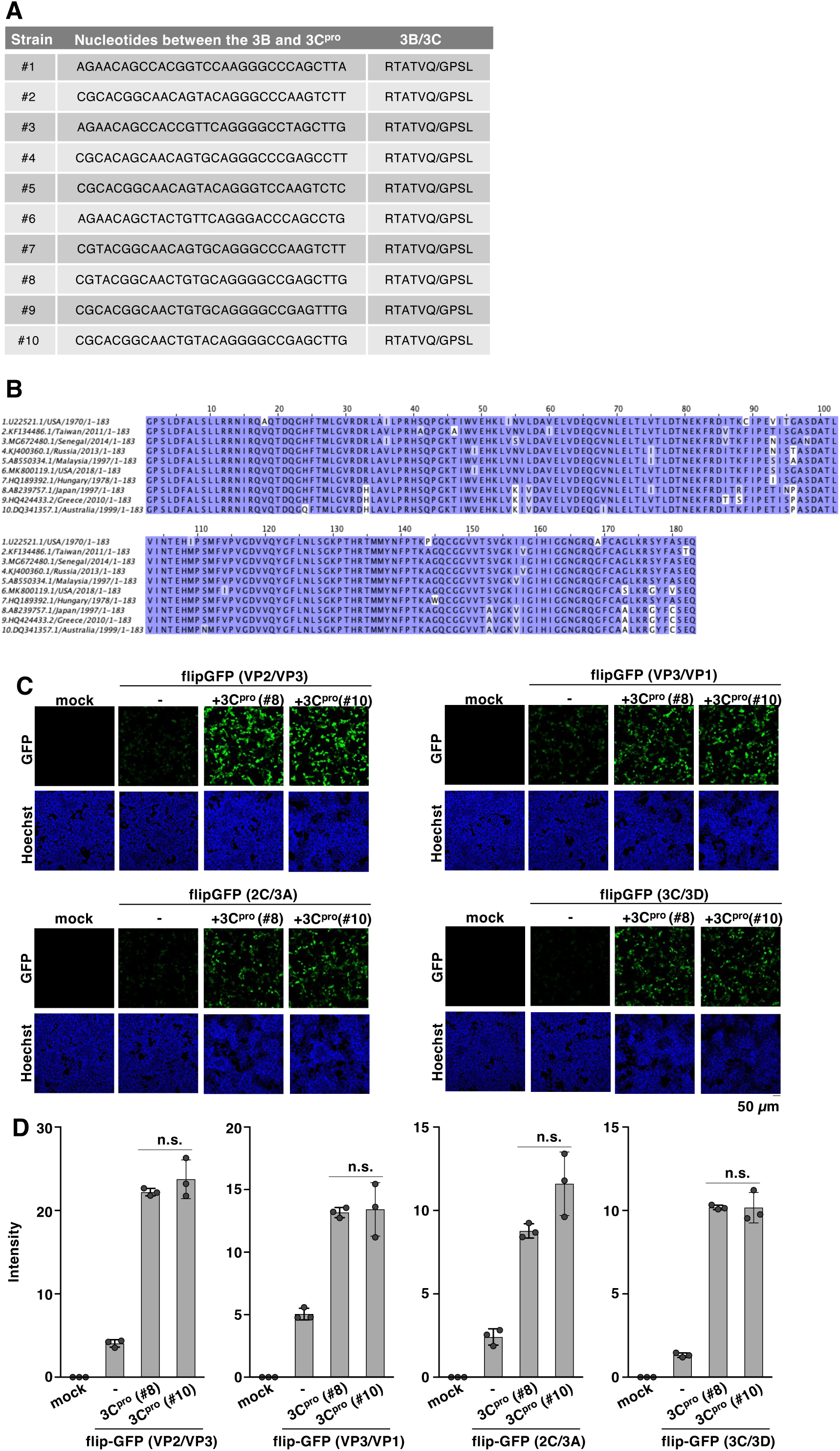
(A) The examined nucleotide and amino acid sequence between the 3B and 3C regions of EV-A71. (B) The amino acid sequence of EV-A71 3C protease used in this study was aligned by Clustal Omega (EMBL-EBI) and indicated using Jalview software version 2.11.4.1 (62). (C) HEK293T cells were transfected with the plasmid encoding the 3C protease of strain #8 or #10 and the flipGFP inserted sequence derived from the VP2/VP3, VP3/VP1, 2C/3A, or 3C/3D sequence of EV-A71. The fluorescent signals were detected using microscopy at 24 h post-transfection. (D) HEK293T cells were transfected with the plasmid encoding the 3C protease of strain #8 or #10 and the flipGFP inserted sequence derived from the VP2/VP3, VP3/VP1, 2C/3A, or 3C/3D sequence of EV-A71. The fluorescent intensity was calculated 24 h post-transfection. The data presented in D is representative of two independent experiments. For the experiment presented in E, significance (n.s., not significant) was determined using Student’s *t-*test (n = 3).

**Supplementary Figure 4.**
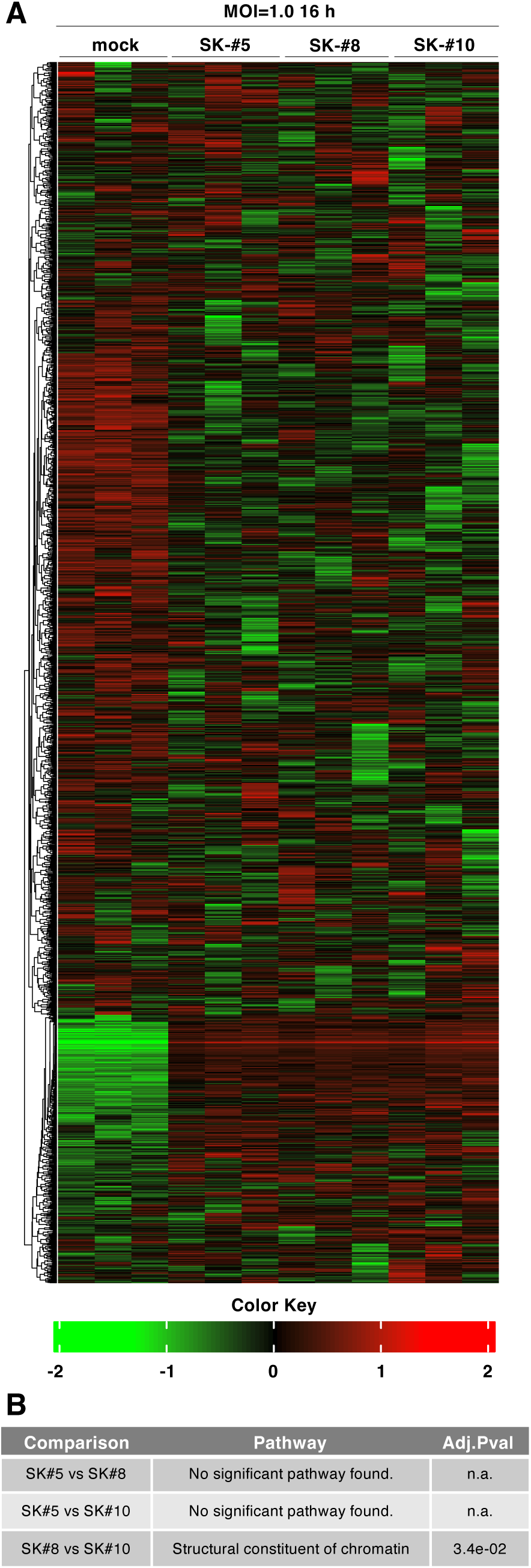
(A) RD-S cells were infected with EV-A71 carrying 3C protease of wild type (SK-#5), strain #8 (SK- #8) or strain #10(SK-#10) at an MOI of 1.0, extracted total RNA at 16 h post-infection and analyzed using RNA sequencing (RNAseq). The result of RNAseq was visualized using a heatmap. (B) Gene ontology (GO) molecular function pathways of RD-S cells infected with EV-A71 were compared using Integrated Differential Expression and Pathway (iDEP) analysis. Adj.Pval, adjusted p-value.

## References

1. Pallansch A, Roos P. 2001. Enteroviruses: Polioviruses, Coxsackieviruses, Echoviruses, and Newer Enteroviruses. Fields Virology 1:pp 723–775.

2. Tuthill TJ, Groppelli E, Hogle JM, Rowlands DJ. 2010. Picornaviruses. Curr Top Microbiol Immunol 343:43–89.

3. Whitton JL, Cornell CT, Feuer R. 2005. Host and virus determinants of picornavirus pathogenesis and tropism. Nat Rev Microbiol 3:765–76.

4. Wright PF, Wieland-Alter W, Ilyushina NA, Hoen AG, Arita M, Boesch AW, Ackerman ME, van der Avoort H, Oberste MS, Pallansch MA, Burton AH, Jaffar MA, Sutter RW. 2014. Intestinal immunity is a determinant of clearance of poliovirus after oral vaccination. J Infect Dis 209:1628–34.

5. Sutter RW, Eisenhawer M, Molodecky NA, Verma H, Okayasu H. 2024. Inactivated Poliovirus Vaccine: Recent Developments and the Tortuous Path to Global Acceptance. Pathogens 13.

6. Yeh MT, Smith M, Carlyle S, Konopka-Anstadt JL, Burns CC, Konz J, Andino R, Macadam A. 2023. Genetic stabilization of attenuated oral vaccines against poliovirus types 1 and 3. Nature 619:135–142.

7. Yeh MT, Bujaki E, Dolan PT, Smith M, Wahid R, Konz J, Weiner AJ, Bandyopadhyay AS, Van Damme P, De Coster I, Revets H, Macadam A, Andino R. 2020. Engineering the Live-Attenuated Polio Vaccine to Prevent Reversion to Virulence. Cell Host Microbe 27:736–751.e8.

8. Lau SK, Woo PC, Lai KK, Huang Y, Yip CC, Shek CT, Lee P, Lam CS, Chan KH, Yuen KY. 2011. Complete genome analysis of three novel picornaviruses from diverse bat species. J Virol 85:8819–28.

9. Du J, Lu L, Liu F, Su H, Dong J, Sun L, Zhu Y, Ren X, Yang F, Guo F, Liu Q, Wu Z, Jin Q. 2016. Distribution and characteristics of rodent picornaviruses in China. Sci Rep 6:34381.

10. Oberste MS, Maher K, Pallansch MA. 2007. Complete genome sequences for nine simian enteroviruses. J Gen Virol 88:3360–3372.

11. Lim ES, Cao S, Holtz LR, Antonio M, Stine OC, Wang D. 2014. Discovery of rosavirus 2, a novel variant of a rodent-associated picornavirus, in children from The Gambia. Virology 454- 455:25–33.

12. Lau SK, Woo PC, Li KS, Zhang HJ, Fan RY, Zhang AJ, Chan BC, Lam CS, Yip CC, Yuen MC, Chan KH, Chen ZW, Yuen KY. 2016. Identification of Novel Rosavirus Species That Infects Diverse Rodent Species and Causes Multisystemic Dissemination in Mouse Model. PLoS Pathog 12:e1005911.

13. Huang SW, Cheng D, Wang JR. 2019. Enterovirus A71: virulence, antigenicity, and genetic evolution over the years. J Biomed Sci 26:81.

14. Shimizu H, Utama A, Yoshii K, Yoshida H, Yoneyama T, Sinniah M, Yusof MA, Okuno Y, Okabe N, Shih SR, Chen HY, Wang GR, Kao CL, Chang KS, Miyamura T, Hagiwara A. 1999. Enterovirus 71 from fatal and nonfatal cases of hand, foot and mouth disease epidemics in Malaysia, Japan and Taiwan in 1997-1998. Jpn J Infect Dis 52:12–5.

15. Baggen J, Thibaut HJ, Strating J, van Kuppeveld FJM. 2018. The life cycle of non-polio enteroviruses and how to target it. Nat Rev Microbiol 16:368–381.

16. Yi J, Peng J, Yang W, Zhu G, Ren J, Li D, Zheng H. 2021. Picornavirus 3C - a protease ensuring virus replication and subverting host responses. J Cell Sci 134.

17. Laitinen OH, Svedin E, Kapell S, Nurminen A, Hytönen VP, Flodström-Tullberg M. 2016. Enteroviral proteases: structure, host interactions and pathogenicity. Rev Med Virol 26:251–67.

18. Weng KF, Li ML, Hung CT, Shih SR. 2009. Enterovirus 71 3C protease cleaves a novel target CstF-64 and inhibits cellular polyadenylation. PLoS Pathog 5:e1000593.

19. de Breyne S, Bonderoff JM, Chumakov KM, Lloyd RE, Hellen CU. 2008. Cleavage of eukaryotic initiation factor eIF5B by enterovirus 3C proteases. Virology 378:118–22.

20. White JP, Cardenas AM, Marissen WE, Lloyd RE. 2007. Inhibition of cytoplasmic mRNA stress granule formation by a viral proteinase. Cell Host Microbe 2:295–305.

21. Tsu BV, Beierschmitt C, Ryan AP, Agarwal R, Mitchell PS, Daugherty MD. 2021. Diverse viral proteases activate the NLRP1 inflammasome. Elife 10.

22. Robinson KS, Teo DET, Tan KS, Toh GA, Ong HH, Lim CK, Lay K, Au BV, Lew TS, Chu JJH, Chow VTK, Wang Y, Zhong FL, Reversade B. 2020. Enteroviral 3C protease activates the human NLRP1 inflammasome in airway epithelia. Science 370.

23. Clark ME, Lieberman PM, Berk AJ, Dasgupta A. 1993. Direct cleavage of human TATA-binding protein by poliovirus protease 3C in vivo and in vitro. Mol Cell Biol 13:1232–7.

24. Das S, Dasgupta A. 1993. Identification of the cleavage site and determinants required for poliovirus 3CPro-catalyzed cleavage of human TATA-binding transcription factor TBP. J Virol 67:3326–31.

25. Zhang Q, Schepis A, Huang H, Yang J, Ma W, Torra J, Zhang SQ, Yang L, Wu H, Nonell S, Dong Z, Kornberg TB, Coughlin SR, Shu X. 2019. Designing a Green Fluorogenic Protease Reporter by Flipping a Beta Strand of GFP for Imaging Apoptosis in Animals. J Am Chem Soc 141:4526–4530.

26. van der Linden L, Ulferts R, Nabuurs SB, Kusov Y, Liu H, George S, Lacroix C, Goris N, Lefebvre D, Lanke KH, De Clercq K, Hilgenfeld R, Neyts J, van Kuppeveld FJ. 2014. Application of a cell-based protease assay for testing inhibitors of picornavirus 3C proteases. Antiviral Res 103:17–24.

27. Zhang Y, Ke X, Zheng C, Liu Y, Xie L, Zheng Z, Wang H. 2017. Development of a luciferase-based biosensor to assess enterovirus 71 3C protease activity in living cells. Sci Rep 7:10385.

28. Saura M, Zaragoza C, McMillan A, Quick RA, Hohenadl C, Lowenstein JM, Lowenstein CJ. 1999. An antiviral mechanism of nitric oxide: inhibition of a viral protease. Immunity 10:21–8.

29. Schmidt NJ, Lennette EH, Ho HH. 1974. An apparently new enterovirus isolated from patients with disease of the central nervous system. J Infect Dis 129:304–9.

30. Pei J, Grishin NV. 2001. AL2CO: calculation of positional conservation in a protein sequence alignment. Bioinformatics 17:700–12.

31. Froggatt HM, Heaton BE, Heaton NS. 2020. Development of a Fluorescence-Based, High-Throughput SARS-CoV-2 3CL(pro) Reporter Assay. J Virol 94.

32. Ma C, Sacco MD, Xia Z, Lambrinidis G, Townsend JA, Hu Y, Meng X, Szeto T, Ba M, Zhang X, Gongora M, Zhang F, Marty MT, Xiang Y, Kolocouris A, Chen Y, Wang J. 2021. Discovery of SARS-CoV-2 Papain-like Protease Inhibitors through a Combination of High-Throughput Screening and a FlipGFP-Based Reporter Assay. ACS Cent Sci 7:1245–1260.

33. Ma C, Hu Y, Wang Y, Choza J, Wang J. 2022. Drug-Repurposing Screening Identified Tropifexor as a SARS-CoV-2 Papain-like Protease Inhibitor. ACS Infect Dis 8:1022–1030.

34. Ma C, Tan H, Choza J, Wang Y, Wang J. 2022. Validation and invalidation of SARS-CoV-2 main protease inhibitors using the Flip-GFP and Protease-Glo luciferase assays. Acta Pharm Sin B 12:1636–1651.

35. Bao HL, Tu G, Yue Q, Liu P, Zheng HL, Yao XJ. 2023. Design and synthesis of isatin derivatives as effective SARS-CoV-2 3CL protease inhibitors. Chem Biol Drug Des 102:857–869.

36. Tan B, Joyce R, Tan H, Hu Y, Wang J. 2023. SARS-CoV-2 Main Protease Drug Design, Assay Development, and Drug Resistance Studies. Acc Chem Res 56:157–168.

37. Hayashi T, Hirano J, Murakami K, Fujii Y, Kobayashi S, Tanikawa T, Kitamura M, Someya Y. 2025. Identification of FDA-Approved Drugs That Inhibit SARS-CoV-2 and Human Norovirus Replication. Biol Pharm Bull 48:994–1000.

38. Andino R, Rieckhof GE, Baltimore D. 1990. A functional ribonucleoprotein complex forms around the 5’ end of poliovirus RNA. Cell 63:369–80.

39. Zell R, Sidigi K, Bucci E, Stelzner A, Görlach M. 2002. Determinants of the recognition of enteroviral cloverleaf RNA by coxsackievirus B3 proteinase 3C. Rna 8:188–201.

40. Shih SR, Chiang C, Chen TC, Wu CN, Hsu JT, Lee JC, Hwang MJ, Li ML, Chen GW, Ho MS. 2004. Mutations at KFRDI and VGK domains of enterovirus 71 3C protease affect its RNA binding and proteolytic activities. J Biomed Sci 11:239–48.

41. Shimizu H, Utama A, Onnimala N, Li C, Li-Bi Z, Yu-Jie M, Pongsuwanna Y, Miyamura T. 2004. Molecular epidemiology of enterovirus 71 infection in the Western Pacific Region. Pediatr Int 46:231–5.

42. Nagata N, Shimizu H, Ami Y, Tano Y, Harashima A, Suzaki Y, Sato Y, Miyamura T, Sata T, Iwasaki T. 2002. Pyramidal and extrapyramidal involvement in experimental infection of cynomolgus monkeys with enterovirus 71. J Med Virol 67:207–16.

43. McMinn P, Lindsay K, Perera D, Chan HM, Chan KP, Cardosa MJ. 2001. Phylogenetic analysis of enterovirus 71 strains isolated during linked epidemics in Malaysia, Singapore, and Western Australia. J Virol 75:7732–8.

44. Yang P, Mathieu C, Kolaitis RM, Zhang P, Messing J, Yurtsever U, Yang Z, Wu J, Li Y, Pan Q, Yu J, Martin EW, Mittag T, Kim HJ, Taylor JP. 2020. G3BP1 Is a Tunable Switch that Triggers Phase Separation to Assemble Stress Granules. Cell 181:325–345.e28.

45. Yang X, Hu Z, Fan S, Zhang Q, Zhong Y, Guo D, Qin Y, Chen M. 2018. Picornavirus 2A protease regulates stress granule formation to facilitate viral translation. PLoS Pathog 14:e1006901.

46. Visser LJ, Langereis MA, Rabouw HH, Wahedi M, Muntjewerff EM, de Groot RJ, van Kuppeveld FJM. 2019. Essential Role of Enterovirus 2A Protease in Counteracting Stress Granule Formation and the Induction of Type I Interferon. J Virol 93.

47. Sidibé H, Dubinski A, Vande Velde C. 2021. The multi-functional RNA-binding protein G3BP1 and its potential implication in neurodegenerative disease. J Neurochem 157:944–962.

48. Mondal S, Sarvari G, Boehr DD. 2023. Picornavirus 3C Proteins Intervene in Host Cell Processes through Proteolysis and Interactions with RNA. Viruses 15.

49. Leonard RA, Rao VN, Bartlett A, Froggatt HM, Luftig MA, Heaton BE, Heaton NS. 2023. A low-background, fluorescent assay to evaluate inhibitors of diverse viral proteases. J Virol 97:e0059723.

50. Tan CW, Tee HK, Lee MH, Sam IC, Chan YF. 2016. Enterovirus A71 DNA-Launched Infectious Clone as a Robust Reverse Genetic Tool. PLoS One 11:e0162771.

51. Hirano J, Hayashi T, Kitamura K, Nishimura Y, Shimizu H, Okamoto T, Okada K, Uemura K, Yeh MT, Ono C, Taguwa S, Muramatsu M, Matsuura Y. 2024. Enterovirus 3A protein disrupts endoplasmic reticulum homeostasis through interaction with GBF1. J Virol 98:e0081324.

52. Niwa H, Yamamura K, Miyazaki J. 1991. Efficient selection for high-expression transfectants with a novel eukaryotic vector. Gene 108:193–9.

53. Yamayoshi S, Yamashita Y, Li J, Hanagata N, Minowa T, Takemura T, Koike S. 2009. Scavenger receptor B2 is a cellular receptor for enterovirus 71. Nat Med 15:798–801.

54. Nishimura Y, Sato K, Koyanagi Y, Wakita T, Muramatsu M, Shimizu H, Bergelson JM, Arita M. 2024. Enterovirus A71 does not meet the uncoating receptor SCARB2 at the cell surface. PLoS Pathog 20:e1012022.

55. Schneider TD, Stephens RM. 1990. Sequence logos: a new way to display consensus sequences. Nucleic Acids Res 18:6097–100.

56. Crooks GE, Hon G, Chandonia JM, Brenner SE. 2004. WebLogo: a sequence logo generator. Genome Res 14:1188–90.

57. Meng EC, Goddard TD, Pettersen EF, Couch GS, Pearson ZJ, Morris JH, Ferrin TE. 2023. UCSF ChimeraX: Tools for structure building and analysis. Protein Sci 32:e4792.

58. Pettersen EF, Goddard TD, Huang CC, Meng EC, Couch GS, Croll TI, Morris JH, Ferrin TE. 2021. UCSF ChimeraX: Structure visualization for researchers, educators, and developers. Protein Sci 30:70–82.

59. Goddard TD, Huang CC, Meng EC, Pettersen EF, Couch GS, Morris JH, Ferrin TE. 2018. UCSF ChimeraX: Meeting modern challenges in visualization and analysis. Protein Sci 27:14–25.

60. Ge SX, Son EW, Yao R. 2018. iDEP: an integrated web application for differential expression and pathway analysis of RNA-Seq data. BMC Bioinformatics 19:534.

61. Ciccarelli FD, Doerks T, von Mering C, Creevey CJ, Snel B, Bork P. 2006. Toward automatic reconstruction of a highly resolved tree of life. Science 311:1283–7.

62. Waterhouse AM, Procter JB, Martin DM, Clamp M, Barton GJ. 2009. Jalview Version 2--a multiple sequence alignment editor and analysis workbench. Bioinformatics 25:1189–91.

